# Phase Separation Can Increase Enzyme Activity by Concentration and Molecular Organization

**DOI:** 10.1101/2020.09.15.299115

**Authors:** William Peeples, Michael K. Rosen

## Abstract

Biomolecular condensates concentrate macromolecules into discrete cellular foci without an encapsulating membrane. Condensates are often presumed to increase enzymatic reaction rates through increased concentrations of enzymes and substrates (mass action), although this idea has not been widely tested and other mechanisms of modulation are possible. Here we describe a synthetic system where the SUMOylation enzyme cascade is recruited into engineered condensates generated by liquid-liquid phase separation of multidomain scaffolding proteins. SUMOylation rates can be increased up to 36-fold in these droplets compared to the surrounding bulk, depending on substrate K_M_. This dependency produces substantial specificity among different substrates. Analyses of reactions above and below the phase separation threshold lead to a quantitative model in which reactions in condensates are accelerated by mass action and by changes in substrate K_M_, likely due to scaffold-induced molecular organization. Thus, condensates can modulate reaction rates both by concentrating molecules and by physically organizing them.

## Introduction

Biomolecular condensates concentrate proteins and RNA molecules without a surrounding membrane^1,2^. Condensates appear in a wide range of biological contexts, including mRNA storage/degradation^3-6^, T-cell activation^7-9^, and ribosome biogenesis^10-12^. Many condensates appear to form through liquid-liquid phase-separation (LLPS), in which oligomerization mediated by multivalent interactions lowers the solubility of proteins and/or nucleic acids sufficiently to form a second phase^1,2,13^. Where measured, the degree of concentration in this phase ranges from 2- to 150-fold for different constituents^14-17^. Scaffold-like molecules, those that contribute more strongly to formation of the condensate, are typically the most highly concentrated components^14,18^. Client-like molecules, which are recruited into the condensate through interactions with scaffolds but do not contribute strongly to formation of the structure, tend to be less concentrated (with exceptions in cases of high affinity scaffold binding^14,18^). The lack of a surrounding membrane allows rapid exchange of most condensate components with the environment without the need for transporters.

Condensates provide potential mechanisms by which cells can rapidly regulate biochemical processes in a spatially defined manner^1,2,19,20^. Because condensates concentrate enzymes and their potential substrates, the compartments are often invoked to function by accelerating biochemical reactions. Indeed, a number of recent studies have shown that enzymatic rates can be increased by LLPS, including nucleation and assembly of actin filaments and microtubules^7,13,21-25^, RNA autosplicing^26^, production of 3’,5’-cyclic GMP-AMP^27^, carboxylation of ribulose-1,5-bisphosphate^28-31^ and activation of Ras^32^. Given that condensates may selectively concentrate certain enzymes in a pathway and/or certain enzyme/substrate pairs, phase separation has also been invoked as a mechanism to provide specificity in signaling or metabolic networks through accelerating some reactions over others^1,33,34^. Since many substrates often compete for the same enzyme, especially in signaling networks, spatial separation could be used to bias some pathways over others. Such bias has been engineered by clustering a metabolic branchpoint enzyme with only one of its two downstream target enzymes, creating a functional auxotroph for the non-clustered pathway^35^.

Beyond mass action (i.e. the effects of simply concentrating components), phase separation could in principle alter reactions by other mechanisms, for example changing molecular conformations^36-38^ and/or inducing specific molecular organizations^39^, and by the effects of crowding on these properties and molecular diffusion. In recent studies of membrane-associated condensates that control actin filament nucleation and Ras activation, phase separation was found to increase the membrane dwell time of key components, consequently increasing their specific activities^22,32^. Thus, in these systems, phase separation accelerates reactions not only by concentrating molecules, but also by changing the intrinsic activities of those molecules. These complexities, however, have not been generally addressed in three-dimensional droplet systems.

Here we sought to develop a simplified *in vitro* system that models natural condensates, which would enable us to understand how phase separation can alter enzymatic activity. We used the FKBP-rapamycin-FRB system^40,41^ to recruit components of the SUMOylation enzymatic cascade^42^ to polySH3 domain and polyProline-Rich-Motif (polyPRM) scaffold proteins^13^. We then measured SUMOylation activity toward several different substrates at scaffold concentrations above and below the LLPS threshold. Recruiting enzyme and substrate into phase separated droplets can substantially increase reaction rates. This effect is dependent on substrate properties: SUMOylation of high K_M_ substrates is accelerated by LLPS, while SUMOylation of others can be inhibited by LLPS due to a combination of low K_M_ and substrate inhibition, thus affording specificity. Quantitative analyses of reactions in droplets and surrounding bulk solution, and at identical concentrations in the absence of scaffolds, demonstrate that condensates can accelerate reactions by two mechanisms: increased concentration, and molecular organization affording a scaffold-dependent decrease in K_M_. Computationally modeling the combined effects of molecular organization (K_M_) and concentration enables description of the reaction rates of various substrates under diverse conditions. These data indicate that three-dimensional condensates, like two-dimensional condensates, can accelerate reactions by both concentration-dependent and concentration-independent mechanisms, affording switchable control over reaction rates and specificity.

## Results

### Design of an inducible condensate-targeted enzyme cascade

We generated a model condensate based on inducible targeting of the SUMOylation enzymatic cascade (Figure 1A)^43^ into phase separated droplets composed of a polySH3 domain protein (polySH3_3_, with three tandem SH3 domains) and its polyproline-rich-motif (polyPRM_5_, with five tandem PRMs) ligand (Figure 1B)^13^. The cascade attaches the 12 kDa Small Ubiquitin-like Modifier protein, SUMO, to diverse substrates. Many of the SUMOylating and deSUMOylating enzymes are concentrated in PML Nuclear Bodies, and it has been suggested that these condensates may regulate cellular SUMOylation in certain conditions^44^. Here we used a minimal version of the cascade, comprising the E1 (SAE1/2 heterodimer) and E2 (Ubc9) enzymes, substrate(s), and conjugatable SUMO1 (Figure 1A).

**Figure 1.**
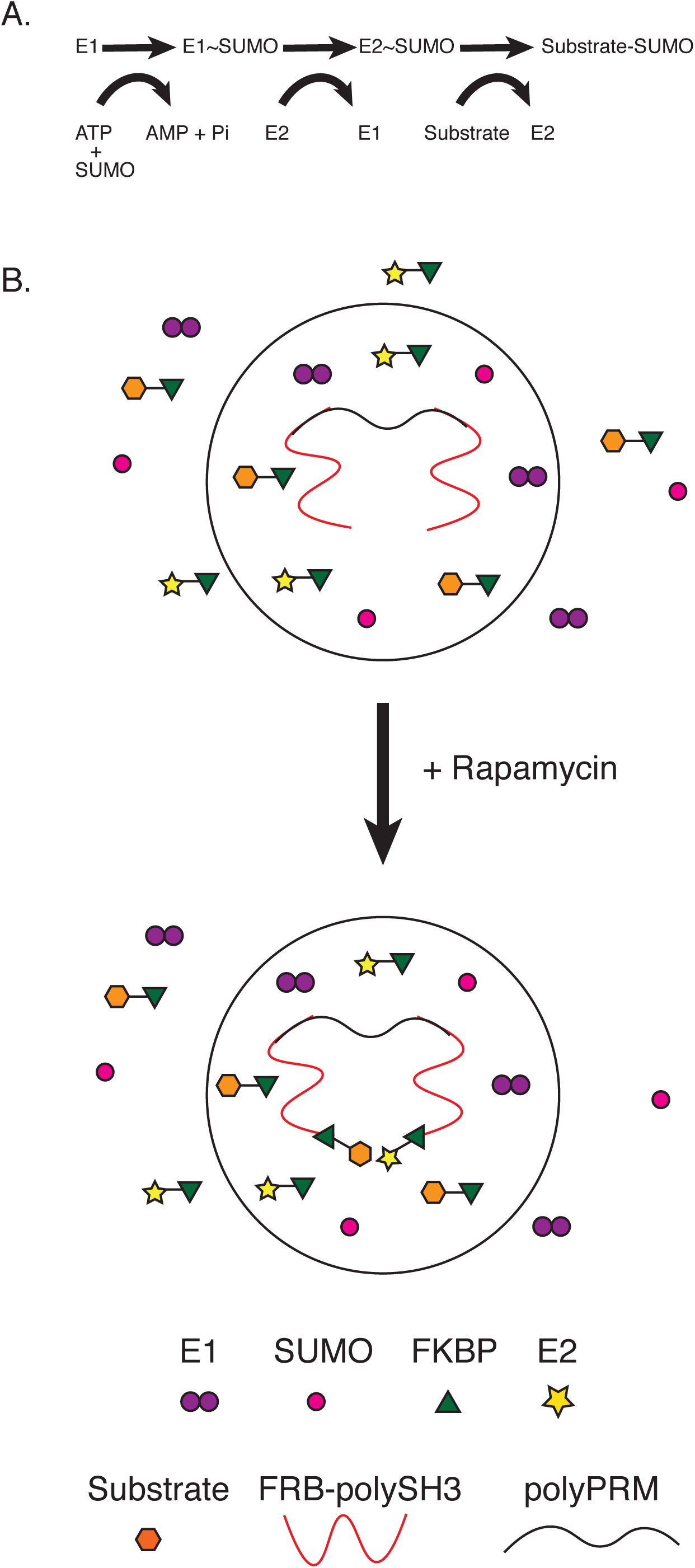
Design of an inducible, condensate-targeted enzyme cascade. (A) Schematic of the SUMOylation cascade. SUMO is initially conjugated to E1 through a thioester bond, then transferred to E2 through a second thioester, and finally to a lysine on the target protein. (B) Inducible recruitment of the SUMOylation cascade to polySH3-polyPRM condensates using the FRB-FKBP-rapamycin system. FRB is fused to polySH3, which is concentrated in the condensates. Upon addition of rapamycin, FKBP-E2 and FKBP-substrate enrich in the condensates, while untagged SUMO remains evenly distributed. Although not illustrated in the diagram, untagged E1 weakly enriches due to binding E2 (see text).

PolySH3_3_ and polyPRM_5_ undergo reversible LLPS when mixed at concentrations above ∼5 µM^13^. We fused the FRB domain from mTOR onto the N-terminus of polySH3_3_, and the FKBP12 protein to the N-termini of Ubc9 and substrates. Addition of rapamycin induces heterodimerization of FRB and FKBP12, recruiting Ubc9 and substrate to the polySH3_3_ scaffold. When polySH3_3_ and polyPRM_5_ are at concentrations below the LLPS threshold they form oligomeric complexes^13^, and rapamycin will cause binding of Ubc9/substrate to these structures. When the scaffolds are above the LLPS threshold, this rapamycin-induced interaction should recruit Ubc9/substrates into the droplet phase (Figure 1C). Note that in all experiments, only E2 and substrate are recruited to the scaffolds by rapamycin, while E1 and SUMO1 are untethered.

### Total SUMOylation activity is increased in the presence of condensates

We initially confirmed that condensates formed by mixing FRB-polySH3_3_ and polyPRM_5_ strongly recruit E2 (mCherry-FKBP-Ubc9) and substrate (FKBP-EGFP-peptide) in the presence of rapamycin but not in a DMSO control (Figure 2A). The substrate used here is a peptide derived from the PML protein (residues 480-495, SQTQSPRKV<u>IKME</u>SEE), which contains the canonical SUMOylation consensus sequence ΨKXE, where Ψ is a large hydrophobic residue, K is the conjugated lysine, X is any amino acid, and E is glutamate^45^. This sequence is recognized directly by Ubc9, bypassing the need for an E3 enzyme.

**Figure 2.**
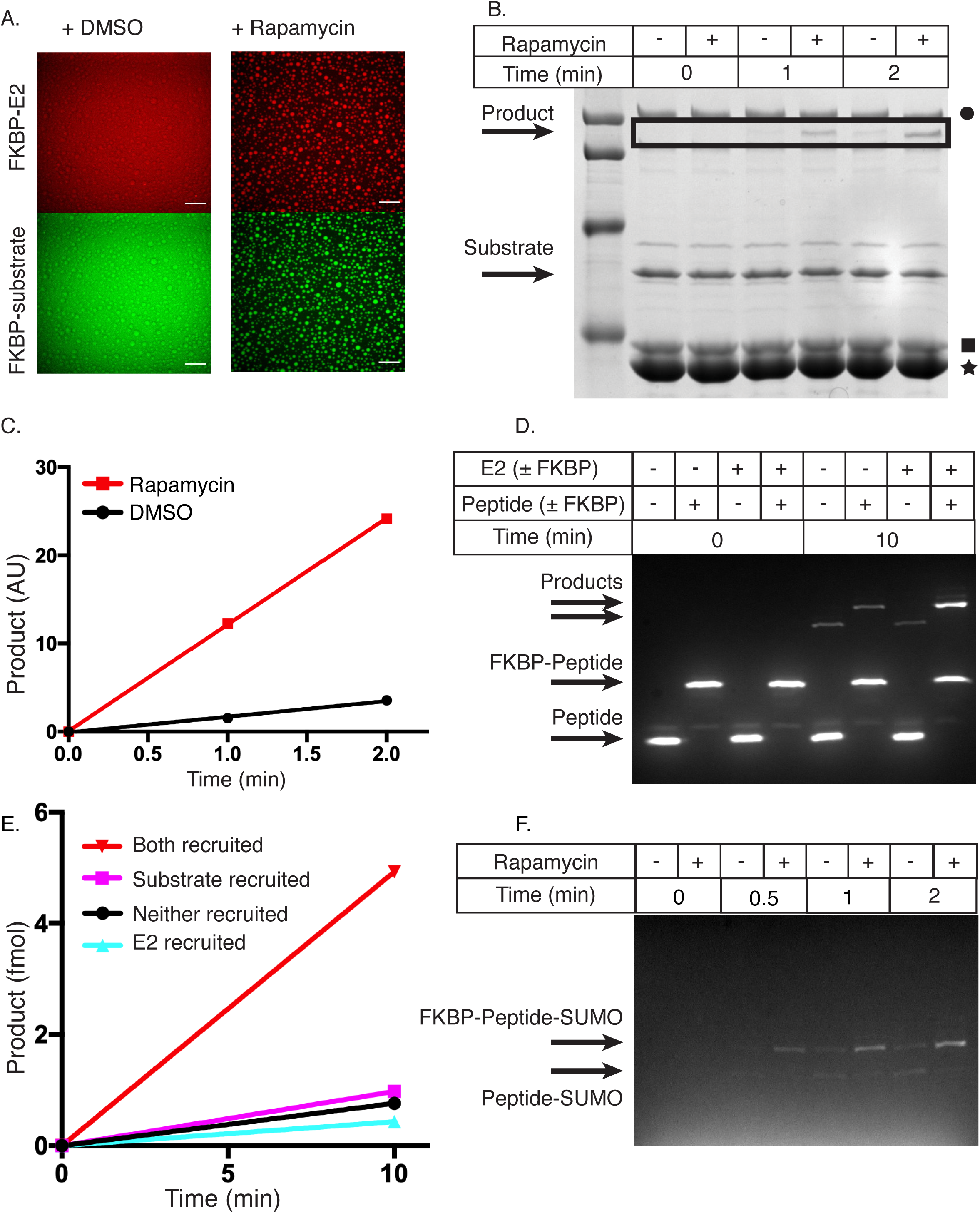
Total SUMOylation activity is increased in the presence of condensates. (A) Confocal fluorescence microscopy images of mCherry-FKBP-E2 (top row) and FKBP-EGFP-substrate (bottom row) in the presence of FRB-polySH3_3_:polyPRM_5_ condensates upon addition of DMSO control (left column) or rapamycin (right column). (B) SDS-PAGE gel stained with Coomassie blue illustrating production of SUMOylated substrate as a function of time with either DMSO (-) or rapamycin (+). Black square denotes E2, black star denotes FRB-polySH3_3_, and black circle denotes E1. (C) Quantification of data in panel B, showing intensity of the SUMOylated substrate band as a function of time. DMSO = black circles, rapamycin = red squares. (D) Fluorescence-detected SDS-PAGE gel depicting production of SUMOylated substrate as a function of time when E2, substrate, both or neither are fused to FKBP (and thus are recruited into FRB-polySH3_3_-polyPRM_5_ droplets). + indicates FKBP-fusion, -indicates non-fused. (E) Quantification of data in panel D, showing intensity of the SUMOylated substrate band as a function of time. E2+substrate recruited, inverted red triangles; E2 recruited, cyan triangles; substrate recruited, magenta squares; neither recruited black circles. Absolute amounts quantified by an internal standard. (F) Fluorescent SDS page gel of SUMOylation of PML peptide or FKBP-PML peptide when co-incubated with FRB-polySH3_3_:polyPRM_5_ condensates and either DMSO or rapamycin. SUMOylated FKBP-peptide is the upper band, and SUMOylated peptide is the lower band.

Even though E1 is not tagged with FKBP, it was moderately enriched in the presence of rapamycin (Figure S1A), likely due to direct interaction with E2, since enrichment requires E2 recruitment (Figures S1A, B). To simplify the microscopy and kinetics of our system, all subsequent experiments used E1 at a saturating concentration, such that the E2∼SUMO1 transfer to substrate is rate-limiting in the cascade, minimizing effects of E1 enrichment (Figure S1C). SUMO1 was not enriched in the droplets in the absence or presence of rapamycin (Figure S1D).

We utilized a gel shift assay to determine whether rapamycin-induced enrichment into droplets modulates activity of the SUMOylation cascade. We mixed all components (E1, mCherry-FKBP-E2, FKBP-EGFP-peptide, SUMO1, FRB-polySH3_3_ and polyPRM_5_) with either DMSO or rapamycin, incubated for 1 hour, and initiated the reaction with ATP. Order of addition did not change the results, suggesting the system equilibrated during the incubation period (not shown). SDS-PAGE analysis showed that the production of SUMO-conjugated product and depletion of FKBP-EGFP-peptide and SUMO1 substrates were much faster with rapamycin than with DMSO control (Figures 2B and C). Acceleration required both FRB-polySH3_3_ and polyPRM_5_ scaffolds and also rapamycin (Figure S2A). Recruitment to FRB-polySH3_3_ alone or the presence of droplets without recruitment does not change the reaction rate. Moreover, acceleration required that both E2 and substrate be recruited together; recruiting only one or the other had little effect (Figures 2D and E). Together these data indicate that the SUMOylation cascade is enhanced by rapamycin-dependent recruitment of E2 and substrate into droplets composed of the FRB-polySH3_3_:polyPRM_5_ complex.

One proposed activity of condensates is to enhance reaction specificity through selective substrate and/or enzyme enrichment. The selective enrichment of FKBP-EGFP-peptide over EGFP-peptide provides a means to test specificity directly in a competitive reaction. We co-incubated EGFP-peptide and FKBP-EGFP-peptide in a single reaction with all other components of the SUMOylation cascade and either DMSO or rapamycin. When incubated with DMSO (no droplet recruitment), both substrates were SUMOylated equally. In the presence of rapamycin, however, FKBP-EGFP-peptide is SUMOylated much more efficiently due to its selective recruitment to the scaffold and into the droplets (Figure 2F). Thus, specificity among otherwise equally reactive substrates can be achieved by selective recruitment into a condensate.

These observations suggest that our condensates enhance total SUMOylation activity when both enzyme and substrate are recruited and can generate specificity via selective substrate recruitment.

### Activity enhancement is substrate-dependent

To learn whether the SUMOylation enhancement we observed with the PML peptide substrate is general, we examined a second substrate, the C-terminal domain of human RanGAP (residues 398-587, hereafter referred to as RanGAP). Unlike the peptide, in addition to its SUMOylation motif (ΨKXE), RanGAP also binds with high affinity to a surface of Ubc9 adjacent to the catalytic site^46^. This interaction positions the RanGAP reactive lysine for conjugation, leading to very efficient SUMOylation with a relatively low K_M_ among substrates (300 ± 200 nM)^47^.

We fused FKBP to RanGAP and recruited it to condensates along with the E2. In contrast to the peptide substrate, SUMOylation of RanGAP is inhibited when the protein is recruited into condensates (Figure 3A and B). This effect is likely significantly due to substrate inhibition at the concentrations achieved in the condensates (Figures S3A, B). An additional factor that could also contribute to decreased total activity upon recruitment is the combination of depletion of substrate from the bulk solution (decreasing enzymatic rate there) coupled with concentration in the droplets to well above K_M_ (which would saturate the E2, producing no compensatory increase in activity there). In general, when K_M_ is low relative to substrate concentration in the droplets, condensates may provide little enhancement in total reaction rate and essentially inhibit activity by sequestration (see Discussion). Consistent with the lack of enhancement being due to one or both of these factors, inhibition was reduced when the assay was performed with a 10-fold lower total concentration of RanGAP (Figure S2B).

**Figure 3.**
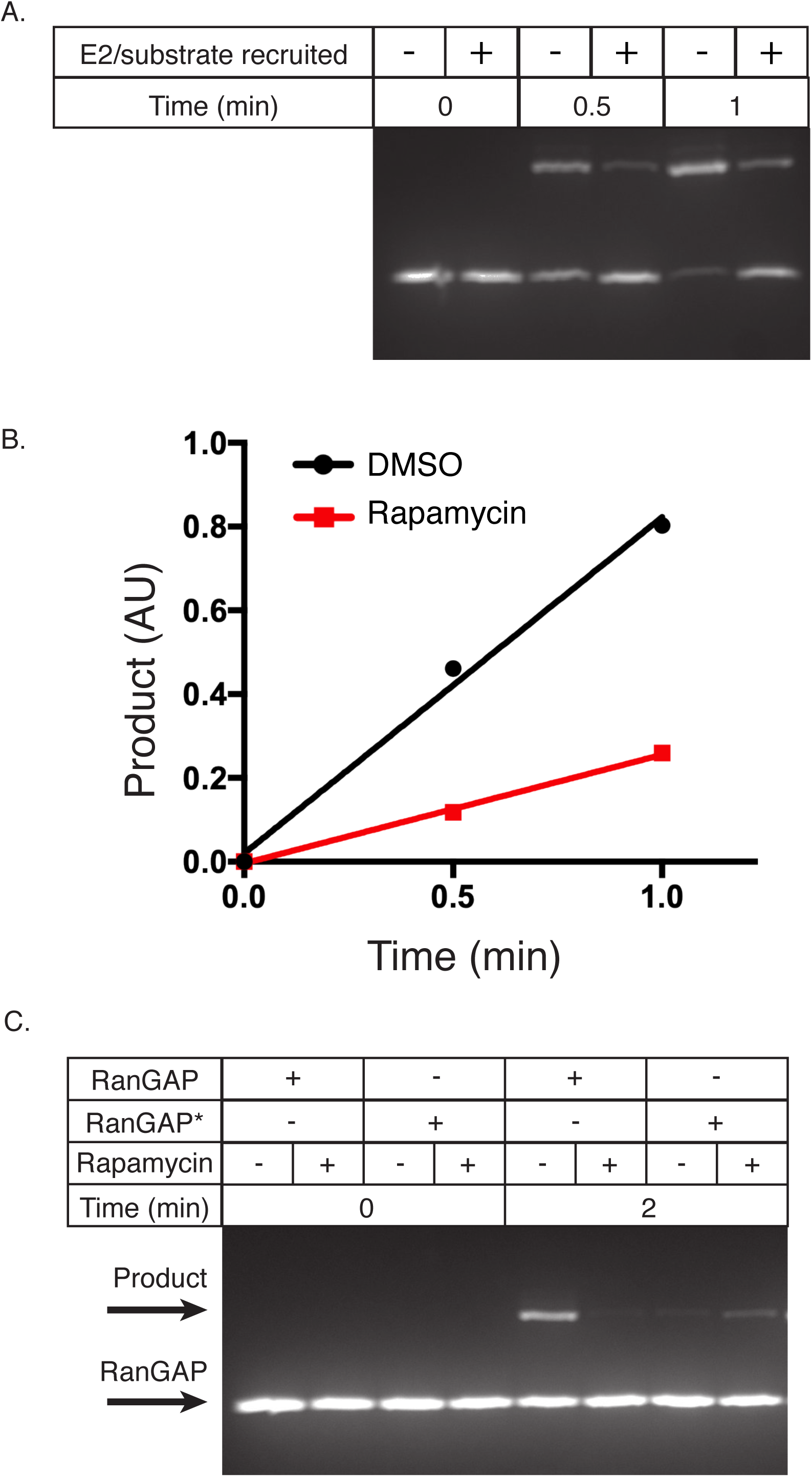
Activity enhancement is substrate-dependent. (A) Fluorescent SDS-PAGE gel depicting the production of SUMOylated substrate as a function of time with FKBP-RanGAP, FKBP-E2, and FRB-polySH3_3_:polyPRM_5_ condensates with DMSO (-) or rapamycin (+). (B) Quantification of data in panel A, showing intensity of the SUMOylated substrate band as a function of time. DMSO, black circles; rapamycin, red squares. (C) Fluorescent SDS-PAGE gel depicting the production of SUMOylated substrate as a function of time with FKBP-RanGAP or FKBP-RanGAP*, FKBP-E2, and FRB-polySH3_3_:polyPRM_5_ condensates with DMSO (-) or rapamycin (+).

To produce a higher K_M_, similar to that of the peptide substrate (∼21 ± 3 µM, Figure S3C), we mutated phenylalanine 562 to alanine in the Ubc9-docking surface of RanGAP ^46^(RanGAP*). This mutation increased K_M_ >100-fold (to 70 ± 8 µM) and also eliminated substrate inhibition (Figure S3D). When recruited to condensates along with E2, the activity of this mutant was appreciably enhanced (Figure 3C). Together, these data suggest that activity enhancement is not a universal property of our condensates but is substrate-dependent. Substrates whose total concentration is low relative to K_M_ are enhanced by recruitment with enzymes into condensates, while substrates whose concentration is high relative to K_M_ and/or display substrate inhibition will not be affected and can even be inhibited by such recruitment.

### SUMOylation is greatly accelerated in the droplet phase

We next sought to quantify the reaction rates in the droplet and bulk phases individually. This required measurement of the total reaction rates in droplet and bulk as well as the volume of each phase.

The droplet volume is sufficiently small in our assays (see below) that it was not feasible to measure activity in droplets directly. Thus, we determined droplet activity by the difference between activity in the total phase separated solution and that in the clarified bulk solution (Figure 4A). For each condition we incubated paired solutions containing FRB-polySH3_3_ and polyPRM_5_ (above the phase separation threshold, generating condensates) plus all components of the SUMOylation cascade: E1, mCherry-FKBP-E2, FKBP-EGFP-RanGAP*, SUMO1, and rapamycin. We removed condensates from one solution of each pair by centrifugation, transferred the supernatant to a separate tube for assay, and initiated reactions by addition of ATP (Figure 4A). SUMOylation measured in the clarified solution yields the bulk reaction rate, while the difference between the reaction rates in each pair represents the droplet rate. The clarified supernatants do not contain visible condensates as assessed by confocal fluorescence microscopy (Figure S4). Sub-diffraction condensates still present in the bulk solution would artifactually decrease the measured droplet activity and increase the measured bulk activity.

**Figure 4.**
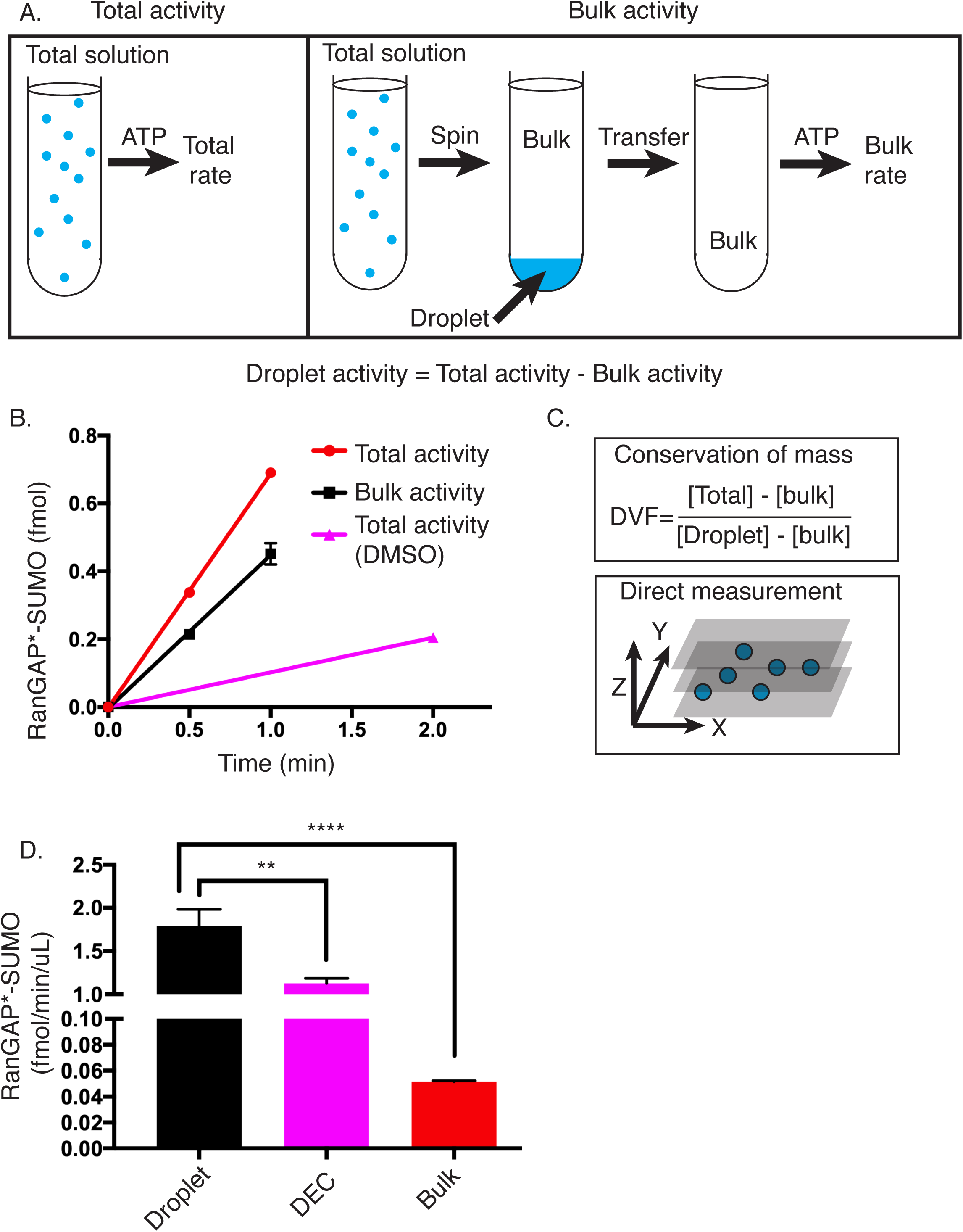
SUMOylation is greatly accelerated in the droplet phase. (A) Schematic of workflow to measure SUMOylation activity in the droplet phase. Total SUMOylation activity is measured by simply mixing all components and adding ATP. Bulk activity is measured by centrifuging the total mixture prior to addition of ATP to sediment droplets, transferring the supernatant to a new tube, and then initiating the reaction with ATP. The difference between Total and Bulk activity yields the Droplet activity. (B) Representative plot showing the production of SUMOylated RanGAP* over time in the total (+rap), bulk (+rap), and total (+DMSO) solutions. Error bars on total (+rap) and bulk (+rap) represent the SEM of 3 experiments. (C) Schematic depicting the two approaches used to calculate droplet volume fraction. Top panel shows the equation used based on conservation of mass. [] represent concentrations in the indicated phase measured by fluorescence imaging. Bottom panel illustrates the direct measurement approach based on confocal imaging of a three-dimensional volume. (D) Volume-normalized activity toward RanGAP* for the droplet, droplet equivalent concentration (DEC), and bulk phases. Error bars represent the SEM from 6 experiments. The statistical significance was assessed by unpaired Student’s t-test. **** represents a p-value < 0.0001.

Addition of rapamycin results in an approximately 7.6-fold increase in total activity in the solution volume (Figure 4B). Analysis of the total and bulk activities revealed that activity is distributed roughly 2:1 between the bulk and droplet compartments. Thus, the reactions are appreciably faster in the droplet phase, since the droplets account for ∼33% of total activity but have much smaller total volume than the bulk.

We next used two independent approaches to determine the droplet volume (Figure 4C). In the first, we initially used calibrated fluorescence intensities to determine the absolute concentrations of enzyme and substrate in the droplet and bulk phases (Table 1) and used conservation of mass to determine their volumes. These analyses revealed that the FRB-polySH3_3_ scaffold is concentrated 105-fold in the droplets. FKBP-E2 and FKBP-RanGAP*, which are both recruited into droplets through rapamycin-mediated tethering to the scaffolds, are concentrated 14-fold and 48-fold, respectively. All of these strongly partitioning components are depleted 10-40 % from the bulk solution (Table 1). Although the E1 enzyme is not tethered to the scaffolds through rapamycin-mediated FKBP-FRB interactions, it nevertheless concentrates in the droplets 1.9-fold through weak binding to E2 (Figure S1A-B). SUMO1 has a partition coefficient of 1, indicating no preference for droplets or bulk solution (Figure S1D). To calculate the volumes of droplet and bulk phases from these measurements, we assumed that the total volume of the mixture is the simple sum of the volumes of solutions added to produce it (i.e. that the volume change upon mixing and LLPS is negligible). With this assumption, the volume fractions of the two phases can be calculated from conservation of mass based on the known total concentration of a given component and its measured concentrations in each compartment (Volume fraction= (total conc – bulk conc)/(droplet conc – bulk conc). For FKBP-EGFP-RanGAP*, this yields a droplet fraction of 1.1± 0.2 %. A similar value of 1.3 ± 0.1 % was found for FRB-polySH3_3_-EGFP, showing consistency for two components with quite different partition coefficients and different mechanisms of droplet enrichment.

As an alternative method, we measured droplet volumes directly. We rapidly imaged a large z-stack through a droplet-containing sample using a spinning disk confocal fluorescence microscope. We joined the images into a three-dimensional volume, identified droplets and summed their volumes. Extrapolation to the total height of the sample and dividing by this volume yielded the volume fraction of the droplets. For droplets containing the complete reaction mixture, and imaged through FKBP-EGFP-RanGAP*, this procedure yielded a droplet volume fraction of 0.9 ± 0.1 %. The similarity of the values determined by the two orthogonal methods lends credence to the approaches and quantitative results. Because of its experimental simplicity and droplet size-independence (see Methods), we used the conservation of mass approach in all analyses below.

Based on the total reaction rates in the droplet and bulk solutions and the known volume fractions, we calculated the reaction rate in each phase, yielding 1.8 fmol/min/µl and 0.05 fmol/min/µl for the droplet and bulk phases, respectively. Thus, under the conditions employed here, the SUMOylation reaction is dramatically accelerated (∼36-fold) within the droplet phase relative to the surrounding bulk phase (Figure 4D).

### Condensates are more active than predicted by concentration alone

We next asked whether the acceleration in condensates is merely due to their higher concentrations of enzymes and substrates, or if there might be additional factors that modulate activity further. To address this question we carried out reactions at the concentrations of E1, E2, RanGAP* and SUMO1 measured within the droplets above (Table 1) but lacking the scaffolds. In these Droplet Equivalent Concentration (DEC) reactions, the SUMOylation rate was 1.1 fmol/min/µl, a value 1.6-fold below that in the droplets (Figure 4D). Thus, while a significant portion of the reaction acceleration produced by condensation derives from the increased concentrations of enzymes and substrates, the higher condensate activity cannot be fully explained by concentration alone. The condensates seem to be imparting excess activity beyond that dictated by mass action.

### Excess activity is driven by a scaffold-induced decrease in apparent K_M_

We considered different mechanisms by which the FRB-polySH3_3_-polyPRM_5_ condensates might impart excess SUMOylation activity. We initially examined macromolecular crowding, which can accelerate reactions by limiting diffusion and/or by excluded volume effects^48^. The combined mass of all scaffolds, enzymes, and substrates in the condensates is ∼30 mg/mL, corresponding to ∼3% w/v. To mimic this condition without scaffolds, we added 3% w/v of either PEG3350 or Ficoll70 to the DEC reaction. Neither agent caused a significant change in activity compared to the DEC conditions alone, suggesting that crowding is not a major contributing factor to excess activity (Figure 5A).

**Figure 5.**
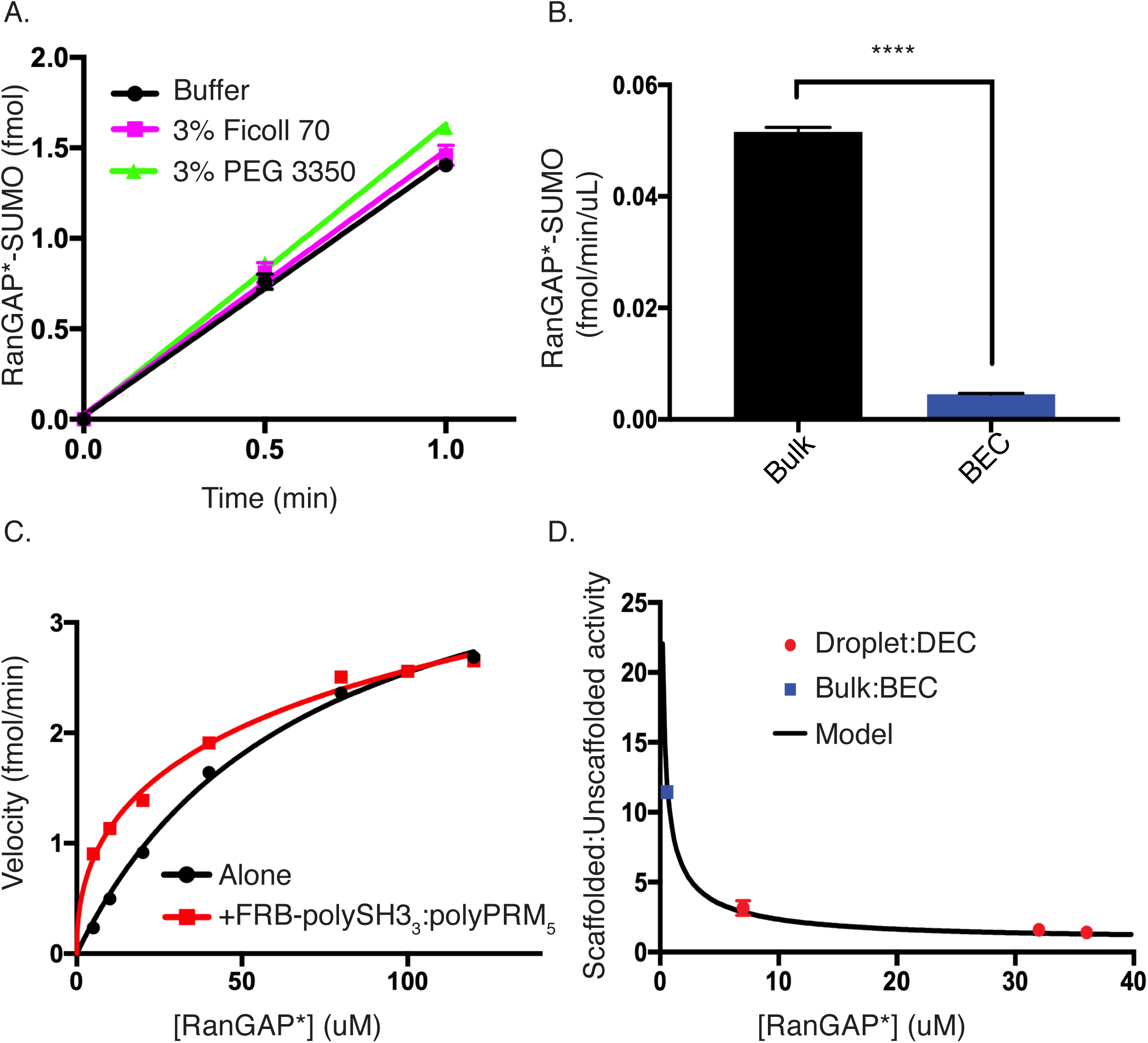
Excess activity is driven by a scaffold-induced decrease in apparent K_M_. (A) Time course showing the production of SUMOylated substrate with E1, E2, RanGAP*, and SUMO1 at droplet equivalent concentrations (black circles), +3% Ficoll 70 (magenta squares), or + 3% PEG 3350 (green triangles). (B) Bar chart showing the volume-normalized activity of bulk (black) or bulk equivalent concentration (BEC, blue). Error bars represent the SEM from 6 experiments. Statistical significance was assessed by unpaired Student’s t-test. **** represents a p-value < 0.0001. (C) Rate of production of SUMOylated RanGAP* as a function of substrate concentration with (red squares) or without (black circles; same data as shown in Figure S3D) sub-threshold concentrations of FRB-polySH3_3_-polyPRM_5_. (D) Ratio of scaffolded:unscaffolded activity (black curve) calculated by dividing the fit of the red curve from the fit of the black curve in Panel C. Overlaid on this curve are the scaffolded:unscaffolded activity ratios from bulk vs BEC (blue square) and droplet vs DEC (red circles). Error bars represent SEM from a total of 3-6 independent measurements.

Another possibility is that FKBP-E2 and FKBP-substrate may be organized by FRB-polySH3_3_:polyPRM_5_ oligomers, analogous to the scaffolding of enzymatic cascades in certain signaling systems^49^. Since FRB-polySH3_3_:polyPRM_5_ oligomers form both above and below the LLPS concentration threshold^13^, we explored this effect by comparing the bulk phase reaction rates with those produced by the same enzyme and substrate concentrations in the absence of scaffolds (Bulk Equivalent Concentration, BEC; Figure 5B). This showed that the scaffolds increase the reaction rate by 11.6-fold (Figure 5B). As in the experiments above on droplets, this enhancement requires both scaffolds and rapamycin-mediated recruitment of both E2 and substrate to them (Figure S5). Thus, the excess activity is not only preserved in the bulk phase, it is even more pronounced there than in the droplets.

To understand the biochemical basis of the scaffold-induced excess activity, we titrated FKBP-EGFP-RanGAP* into the SUMOylation reactions in the presence and absence of sub-LLPS threshold concentrations of FRB-polySH3_3_ and poly-PRM_5_. As shown in Figure 5C, the scaffolds shift the curve to the left, indicating a decrease in apparent K_M_. We attempted to model the scaffolded reaction data with simple Michaelis-Menten kinetics, but the fit was poor (not shown). The data could be fit well, though, to Michaelis-Menten with negative cooperativity (Hill coefficient = 0.46), yielding a K_M_ value of 17 ± 12 µM (versus 70 µM for the unscaffolded reaction), with no change in V_max_. We believe this negative cooperativity may derive from the necessity of having both enzyme and substrate bound simultaneously to proximal sites in a scaffold oligomer in order to enhance the reaction. Such dual binding becomes less frequent at high FKBP-RanGAP* concentrations, effectively returning the system to the unscaffolded state. We note that the scaffolded K_M_ is very likely a weighted average of a distribution of different K_M_ values due to different oligomeric states of the scaffolds and organization of enzyme and substrate attached to them.

One consequence of the shift in K_M_ is that the ratio of scaffolded to unscaffolded activity, ([S]^h^/(K _M,S_ ^h^ + [S]^h^)/([S]/(K _M,US_ + [S]) where h is the Hill coefficient, is largest (=[S]^h-1^*K_M,US_/ K _M,S_ ^h^) at low concentrations and decreases to 1 as the substrate approaches saturation (Figure 5D). This behavior is qualitatively consistent with our observation that excess activity is higher in the low-concentration bulk than in the high-concentration droplets. Thus, we asked whether the excess activity in the droplet phase could also be described by a scaffold-induced change in K_M_ from 70 µM to 17 µM. As shown in Figure 5D, for three different total concentrations of substrate (0.5, 1.0, 2.0 µM), producing droplet concentrations spanning from 7 µM to 36 µM, the excess activity (now defined as the ratio of scaffolded (droplet or bulk) to unscaffolded (DEC or BEC) rates) in the droplets and bulk fall on the same curve. Together, these data suggest that excess activity arises from changes in K_M_ due to organization of E2 and substrate by the oligomeric FRB-polySH3_3_:poly-PRM5 scaffold. This effect is analogous in droplet and bulk, but manifests differently due to differences in substrate concentration relative to K_M_ in the two phases.

### Activity enhancement is scaffold-specific

We next asked whether these effects differ between scaffolds. Our previous data were generated using a trimeric SH3 protein, polySH3_3_. We repeated the substrate titration in the presence of the analogous pentameric SH3 scaffold, FRB-polySH3_5_, and polyPRM_5_ below the LLPS threshold concentration. In contrast to the trimeric scaffold, the pentameric FRB-polySH3 had no effect on reaction rates when combined with polyPRM_5_. Thus, K_M_ and V_max_ are unaffected by the higher valence scaffold (Figure 6A). The lack of an effect is not due to a tethering defect, since condensates formed by polySH3_5_:polyPRM_5_ recruit FKBP-E2 and FKBP-RANGAP* (in the presence of rapamycin) to virtually the same degree as those formed by the trimeric scaffold (1.3 vs 1.4 µM, and 32 vs 31 µM, respectively).

**Figure 6.**
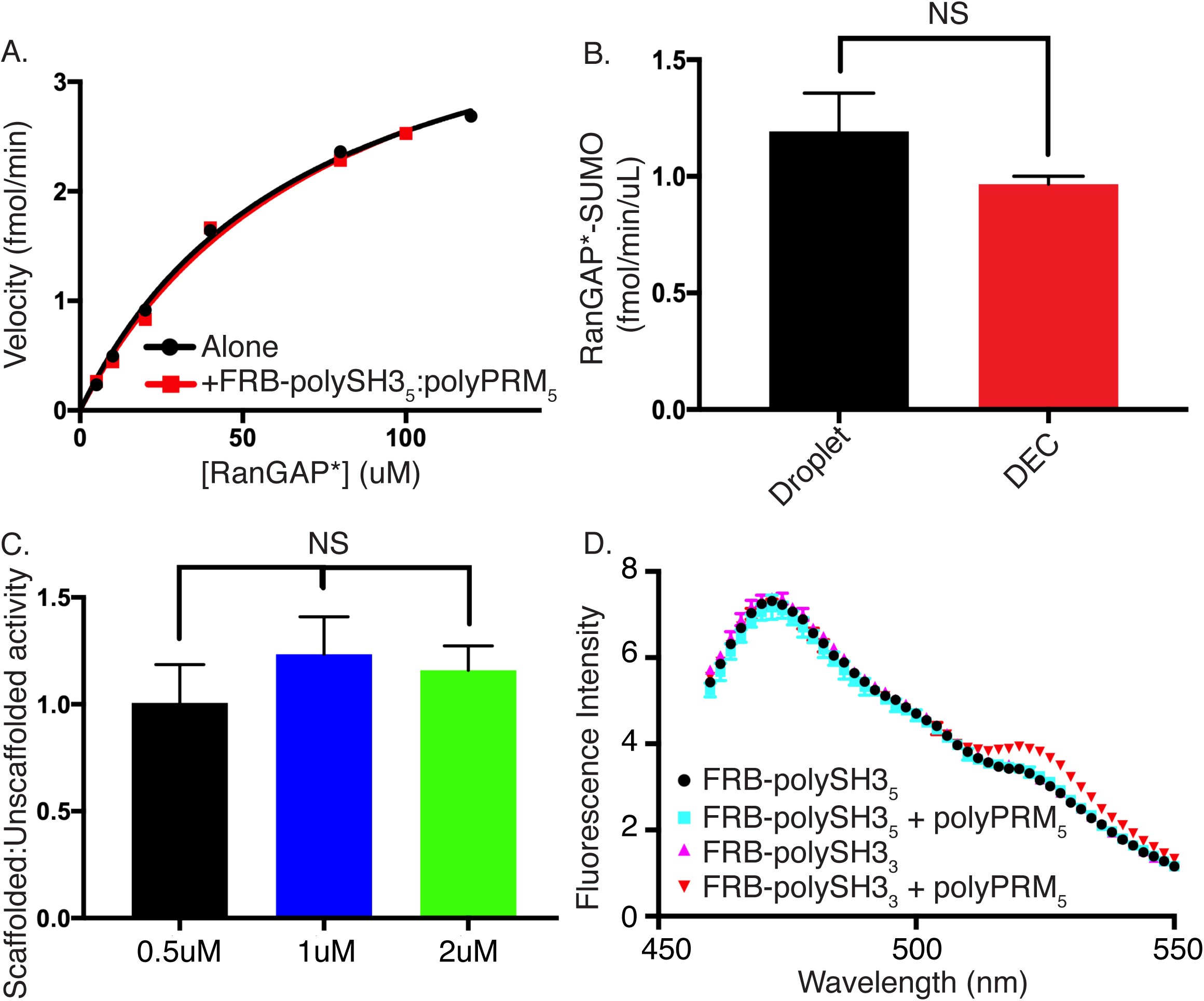
Activity enhancement is scaffold-specific. (A) Rate of production of SUMOylated RanGAP* as a function of RanGAP* concentration with (red) or without (black; same data as shown in Figure S3D) subcritical concentrations of FRB-polySH3_5_-polyPRM_5_. (B) Volume-normalized activity of FRB-polySH3_5_-polyPRM_5_ droplets (black) and droplet equivalent concentration (DEC, red). Error bars represent SEM from 6 independent experiments. Statistical significance was assessed by Student’s t-test with a p-value cutoff of 0.05. (C) Droplet:DEC activity ratio of FRB-polySH3_5_-polyPRM_5_ droplets at total RanGAP* concentrations of 0.5 µM (black), 1 µM (blue), and 2 µM (green), corresponding to droplet concentrations of ∼9 µM, ∼32 µM, and ∼40 µM, respectively. Error bars represent SEM from 6 experiments. Statistical significance was assessed by Student’s t-test with a p-value cutoff of 0.05. (D) Fluorescence emission spectrum of FKBP-YPet-RanGAP* upon 445nm excitation of CyPet-FKBP-E2. Spectra recorded in the presence of FRB-polySH3_5_ (black circles), FRB-polySH3_5_ + polyPRM_5_ (cyan squares), FRB-polySH3_3_ (magenta triangles) or FRB-polySH3_3_ + polyPRM_5_ (inverted red triangles).

We then examined the effects of LLPS by the pentameric scaffold system on the SUMOylation reaction. As with the trimeric scaffold, recruiting FKBP-E2 and FKBP-RANGAP* into the FRB-polySH3_5_:polyPRM_5_ droplets substantially increased the SUMOylation rate. However, using the procedures above to quantitatively compare activity in droplets with that in DEC conditions, we found that the pentameric analog produced no excess activity (Figure 6B). Moreover, doubling or halving the substrate concentration did not produce excess activity (Figure 6C). Thus, for the pentameric scaffold, which does not alter K_M_, the effects of phase separation on SUMOylation rates can be quantitatively described simply by mass action.

Together, these data further support our model in which excess activity induced by the polySH3:polyPRM scaffolds results from tethering-dependent changes in K_M_. Further, because the K_M_ effects are scaffold-specific, they show that LLPS driven by different scaffolds can have different effects on activity, with some systems acting purely through mass action and others acting additionally to change the kinetic parameters of the reaction.

Finally, we asked whether the FRB-polySH3:polyPRM oligomers, which can possess multiple binding sites for FKBP-tagged proteins, could bring E2 and substrate into spatial proximity, which could decrease the apparent K_M_ of the SUMOylation reaction^50^. We tagged FBKP-E2 and FKBP-RanGAP* with CyPet and YPet, respectively, fluorescent proteins that undergo fluorescence resonance energy transfer (FRET) at distances < 100 Å (reference ^51^). We incubated the labeled enzyme and substrate (plus rapamycin) with either the trimeric or pentameric FRB-polySH3 scaffold below the LLPS threshold concentration in the presence or absence of polyPRM_5_ and measured CyPet-YPet FRET. With the pentameric SH3 scaffold, no FRET is observed either with or without polyPRM_5_. In contrast, while the trimeric FRB-polySH3 scaffold does not produce FRET on its own, further addition of polyPRM_5_ produces a reproducible FRET signal (Figure 6D). This signal is dependent on rapamycin, indicating that it results from recruitment of the proteins to the oligomeric scaffold (Figure S6A). These data suggest that the trimeric scaffold brings E2 and substrate closer together on average than the pentameric scaffold, and thus has a greater effect on the SUMOylation reaction.

## Discussion

The engineered system that we have developed here captures important aspects of natural biomolecular condensates. It is composed of a small number of multivalent scaffold molecules that produce the condensate through LLPS, and a larger number of client molecules that are recruited through binding to these scaffolds^18,52^. Recruitment of clients into the condensate can be triggered through environmental factors (here, rapamycin). The degrees of component concentration (partition coefficients) are in the 2 ∼ 100 range, which reflects that observed in condensates where partitioning has been quantified^14,53,54^. Finally, the droplet volume fraction in our system, ∼1 %, is similar to that observed for natural condensates^14^.

Using this system, we have found that recruitment to phase separated scaffolds can enhance enzymatic reactions by up to 36-fold within the condensate compared to the surrounding bulk, and ∼7-fold overall in the total reaction volume. Enhancement in the droplets is achieved through two different mechanisms, mass action and also a scaffold-dependent decrease in K_M_ of the reaction, which results from scaffold-induced molecular organization. The latter effect also accounts for the scaffold-dependent increase in activity of the dilute phase, which also contributes to the increased total activity.

These analyses allow us to quantitatively understand the effects of recruiting the SUMOylation cascade into phase separated droplets under the conditions used here with RanGAP* as substrate. In the absence of rapamycin, the solution has very low activity (0.01 fmol/min/µL), which is evenly distributed in solution. Upon addition of rapamycin, E2 and RanGAP* (and to a much lesser extent, E1) are concentrated in the droplets and depleted from the bulk. Compared to the rate without rapamycin, the rate in the bulk increases due to the reduced K_M,S_ to 0.05 fmol/min/µL. Within the droplets, the rate is 36-fold higher than in the bulk, at 1.8 fmol/min/µL. A droplet volume of 1.1%, thus yields a total activity of 0.07 fmol/min/µL (0.989*0.05 fmol/min/µL + 0.011*1.8 fmol/min/µL; Figure 4B). Addition of rapamycin not only results in an increase in total activity but activity increases in both phases, giving rise to a total activity increase of 7.3-fold. In the absence of rapamycin, the bulk phase contributes 98.9% of the total activity, while the droplet phase contributes 1.1%. In the presence of rapamycin, the bulk phase contributes 70% of total activity, while the droplet phase contributes 30%.

The overall ∼7-fold reaction enhancement that we observed here is relatively modest (although such increases could be significant in biological systems). However, as mentioned above, while E1, E2 and substrate were all enriched in the condensates, only the E2 reaction was enhanced, since we chose conditions that saturated the E1 reaction to simplify the data. In other systems where multiple elements of a pathway are enriched in a condensate and all are functioning at non-saturating conditions, the flux through the cascade should increase by the product of the individual step enhancements. Thus, one can envision quite substantial increases in flux, and also specificity, in multi-step processes. Consistent with this idea, preliminary data on condensate-mediated modification with SUMO2, which unlike SUMO1 can be multiply conjugated to produce polySUMO chains, suggest rate increases upon droplet recruitment much greater than those observed here (not shown). We believe that such multi-step cascades are the most likely beneficiaries of recruitment to condensates *in vivo*, an idea that may explain why many condensates recruit multiple components of complex pathways^55-57^.

To better understand the interplay between partition coefficient, scaffolding effects on K_M_, and substrate concentration, we modeled the change in total reaction rate upon recruitment of enzyme and substrate into condensates. Initially, we assumed a K_M,US_/K_M,S_ ratio of 4.1, as observed here (Figure 5C) and equal substrate and enzyme partition coefficients. Extended Data Figure 1A illustrates how the activity ratio (= total scaffolded reaction / total unscaffolded reaction) varies with substrate concentration (expressed as [S]/K_M,US_) and partition coefficient. For low substrate concentrations, increased partition coefficient produces higher total reaction rate, as enzyme and substrate are concentrated together in the droplets. However, the effect of partition coefficient decreases as substrate concentration grows, with a pronounced shift in behavior near [S] ∼ K_M,US_ (Extended Data Figure 1A, inset), such that when [S] >> K_M,US,_ recruitment of the system into droplets has no effect on the total reaction rate. As a corollary, the sensitivity of the system to partition coefficient decreases as substrate concentration increases. If only substrate has a large partition coefficient, saturation of enzyme within the droplets can cause the total rate to decrease, due to depletion of substrate in the bulk (Figure S8).

We also plotted the activity ratio as a function of partition coefficient and K_M,US_/K_M,S_ at low and high substrate concentrations (Extended Data Figures 1B and C, respectively). At low substrate concentration, increasing K_M,US_/K_M,S_ increases the activity ratio, up to a plateau value when [S] >> K_M,S_. This increase is synergistic with that due to partition coefficient. At high substrate concentration, activity ratio is damped; partition coefficient has no effect, and the K_M_ ratio has a very minor effect that quickly saturates.

Finally, we examined the ratio of droplet to bulk reaction rate as a function of partition coefficient and substrate concentration. This ratio increases strongly with partition coefficient due co-enrichment of enzyme and substrate in the droplets and co-depletion in the bulk. As with the total activity, the ratio is damped at high substrate concentrations as the enzyme becomes saturated (Figure S7A). The fraction of total activity in the droplets parallels these trends as well (Figure S7B).

Our data and modeling illustrate the ability of condensates to impart substrate specificity through several different mechanisms. First, by selectively recruiting some substrates over others, condensates can direct flux through one pathway over another (Figure 2F). In a second mode, substrates at low concentrations relative to their K_M_ values will benefit more from condensate recruitment than those at high concentrations (Extended Data Figure 1A, Figure S7). For substrates such as RanGAP, that possess substrate inhibition, recruitment can be inhibitory (Figure 3), and might serve to sequester the enzyme against other substrates. Finally, we have shown that different scaffolds can impact activity through different modes, with one acting purely through mass action and another through both mass action and a decrease in the apparent K_M_ of the SUMOylation reaction, resulting in excess activity (Figures 5 and 6). Thus, the nature of the scaffold, and likely the way in which individual enzymes and substrates are recruited to it, can impact its effects on activity. In natural systems these mechanisms are expected to act together to impart substantial specificity on biochemical processes occurring within condensates.

In cells the SUMOylation cascade is opposed by SUMO proteases, and the balance of these systems tunes the net level of substrate SUMOylation^58^. Our system has not addressed this additional complexity. Likewise, in PML nuclear bodies the relative enrichment of SUMOylation and deSUMOylation components has not been investigated. In both cases, differential recruitment of the opposing activities could enhance or suppress the impact of condensates on SUMOylation levels and substrate specificity. Additional quantitative biochemical and cellular studies, as well as modeling, will be necessary to understand the full spectrum of behaviors enabled by recruitment of the SUMO regulatory machinery into condensates.

Our model system has provided evidence of sequestration, substrate specificity, and factors beyond mass action in dictating the consequences of enzyme recruitment into condensates. As genomics, proteomics, and imaging improve our understanding of condensate properties, it will be important to understand how factors such as composition and dynamics further modulate these mechanisms to control cellular biochemistry. The ability to quantitatively measure biochemical activities in natural condensates, both reconstituted in vitro and in cells, represents an exciting (and challenging) direction for future studies.

## Acknowledgements

We thank Salman Banani and Allyson Rice for constructs, and all members of the Rosen Lab, past and present, for helpful advice and discussions. Research was supported by the Howard Hughes Medical Institute, a Paul G. Allen Frontiers Distinguished Investigator Award and a grant from the Welch Foundation (I-1544 to M.K.R.).

## Author Contributions

M.K.R. and W.P. conceived the study and designed the research program. W.P. performed all experiments. M.K.R. secured funding and supervised the work. W.P. and M.K.R. wrote the manuscript.

---

**Extended Data Figure 1.**
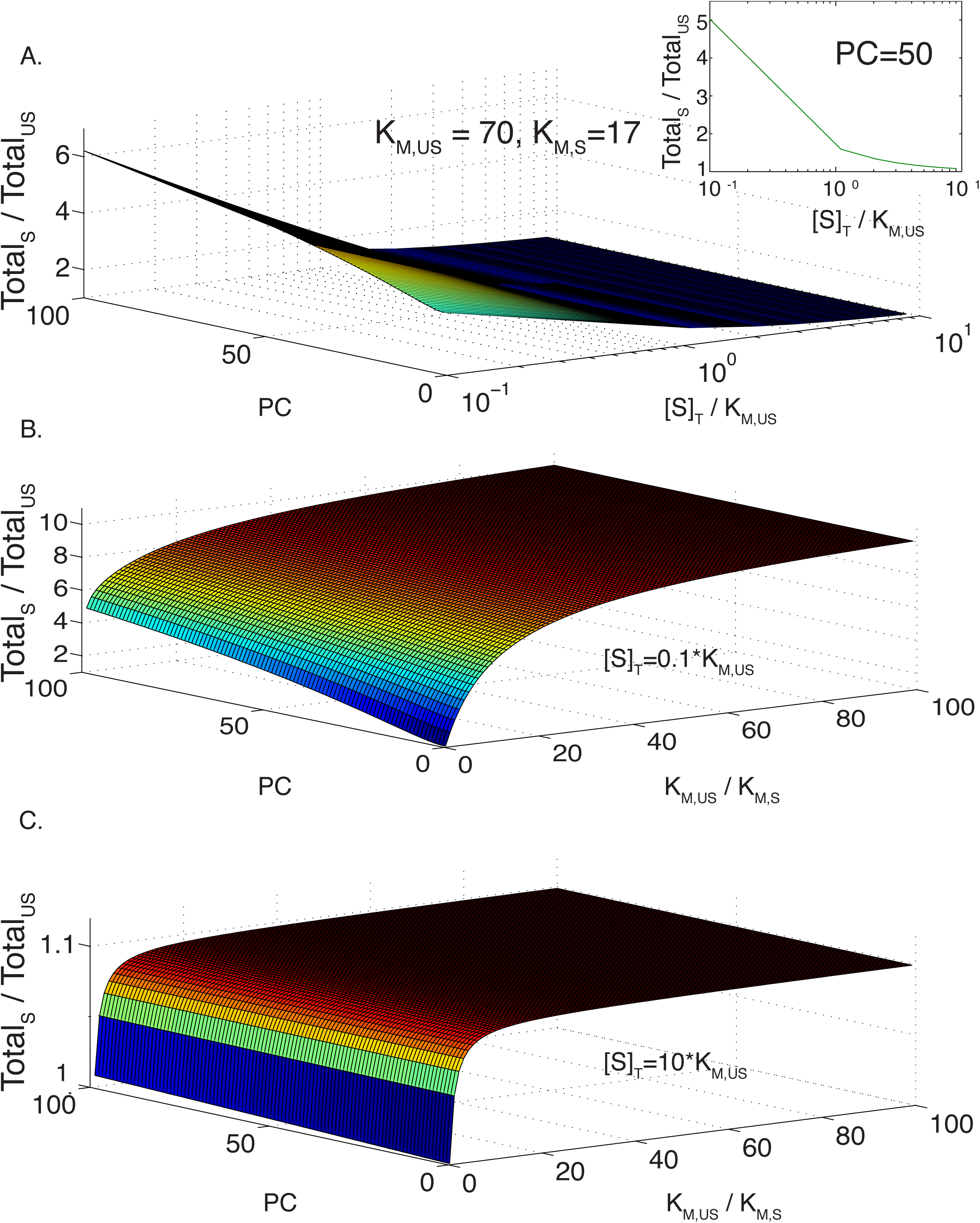
Sensitivity of enhanced condensate activity to K_M_, substrate concentration, and partition coefficient. (A) Modeled ratio of total activity in a phase separated solution, with and without recruitment of enzyme and substrate to the scaffold (Total_S_ and Total_US_, respectively, as a function of substrate concentration (plotted as [S]/K_M,US_) and partition coefficient (PC). Modeled for K_M,US_ = 70 and K_M,S_ = 17, as measured for FRB-polySH3_3_+polyPRM_5_, with identical PC values for enzyme and substrate. Modeling assumes simple, hyperbolic Michaelis-Menten kinetics (see Methods). Color scale is a relative representation of the z-axis values and goes from low (blue) to high (red). Inset is a plot of Total_S_:Total_US_ activity as a function of substrate concentration at a fixed partition coefficient of 50. (B) Modeled ratio of Total_S_ to Total_US_ as a function of PC and the change in K_M_ upon recruitment of enzyme and substrate to the scaffold, K_M,US_/K_M,S_. Total substrate concentration, [S]_T_, set to 0.1 * K_M,US_. (C) Same as (B), except [S]_T_ set to 10 * K_M,US_.

## Supplemental Figure Legends

**Figure S1.**
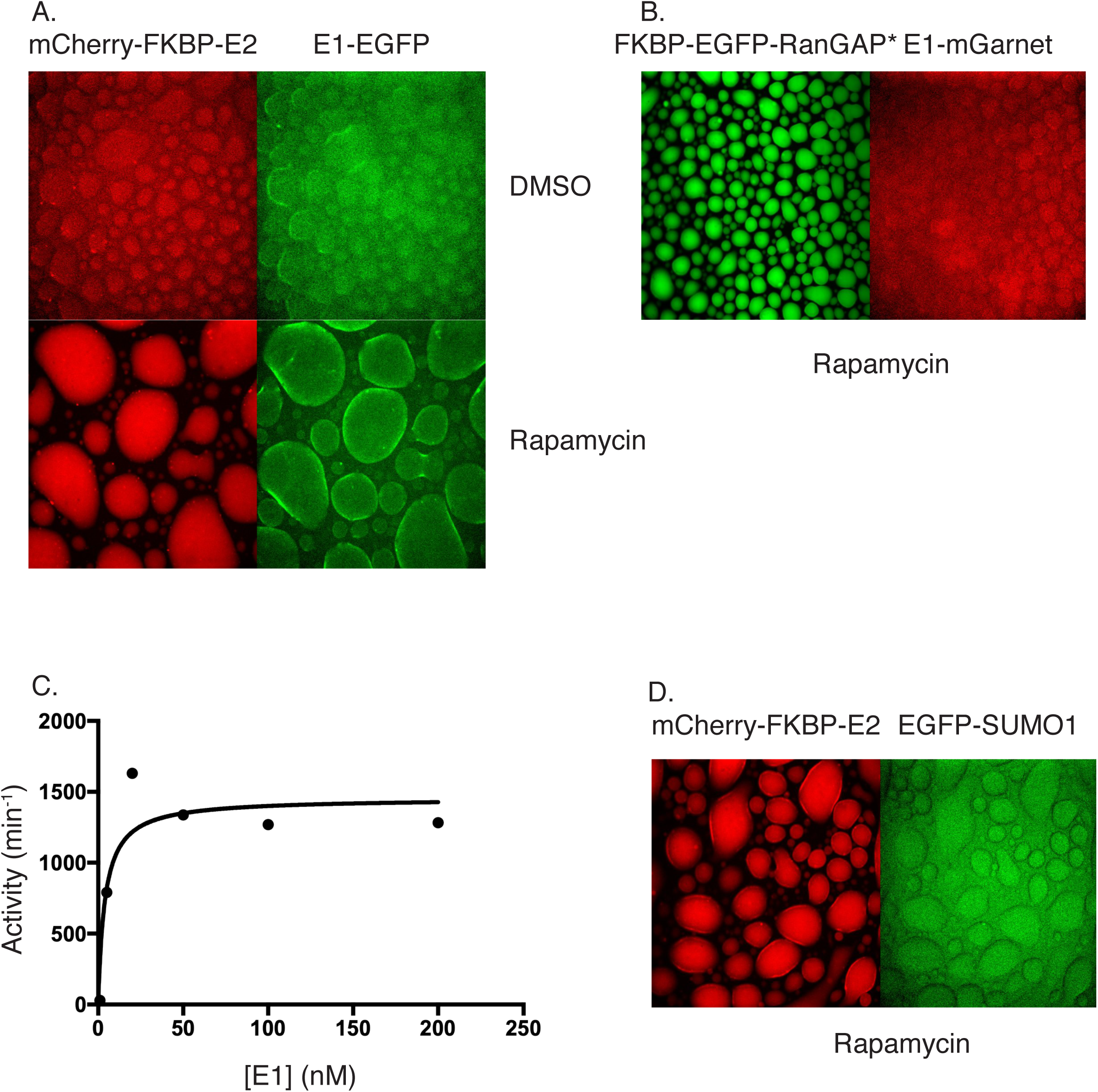
E2-Mediated recruitment of E1, and no recruitment of SUMO. (A) Confocal fluorescence microscopy images of mCherry-FKBP-E2 (red) and E1-EGFP (green) in the presence of FRB-polySH3_3_-polyPRM_5_ condensates with DMSO (top row) and rapamycin (bottom row). (B) Confocal fluorescence microscopy images of FKBP-EGFP-substrate (green) and E1-mGarnet (red) in the presence of FRB-polySH3_3_-polyPRM_5_ condensates and rapamycin. (C) Activity curve depicting SUMOylation of RanGAP* as a function of E1 concentration. Apparent E1 K_M_ is approximately 5 nM. (D) Confocal fluorescence microscopy images of mCherry-FKBP-E2 (red) and EGFP-SUMO1 (green) in the presence of FRB-polySH3_3_-polyPRM_5_ condensates and rapamycin.

**Figure S2.**
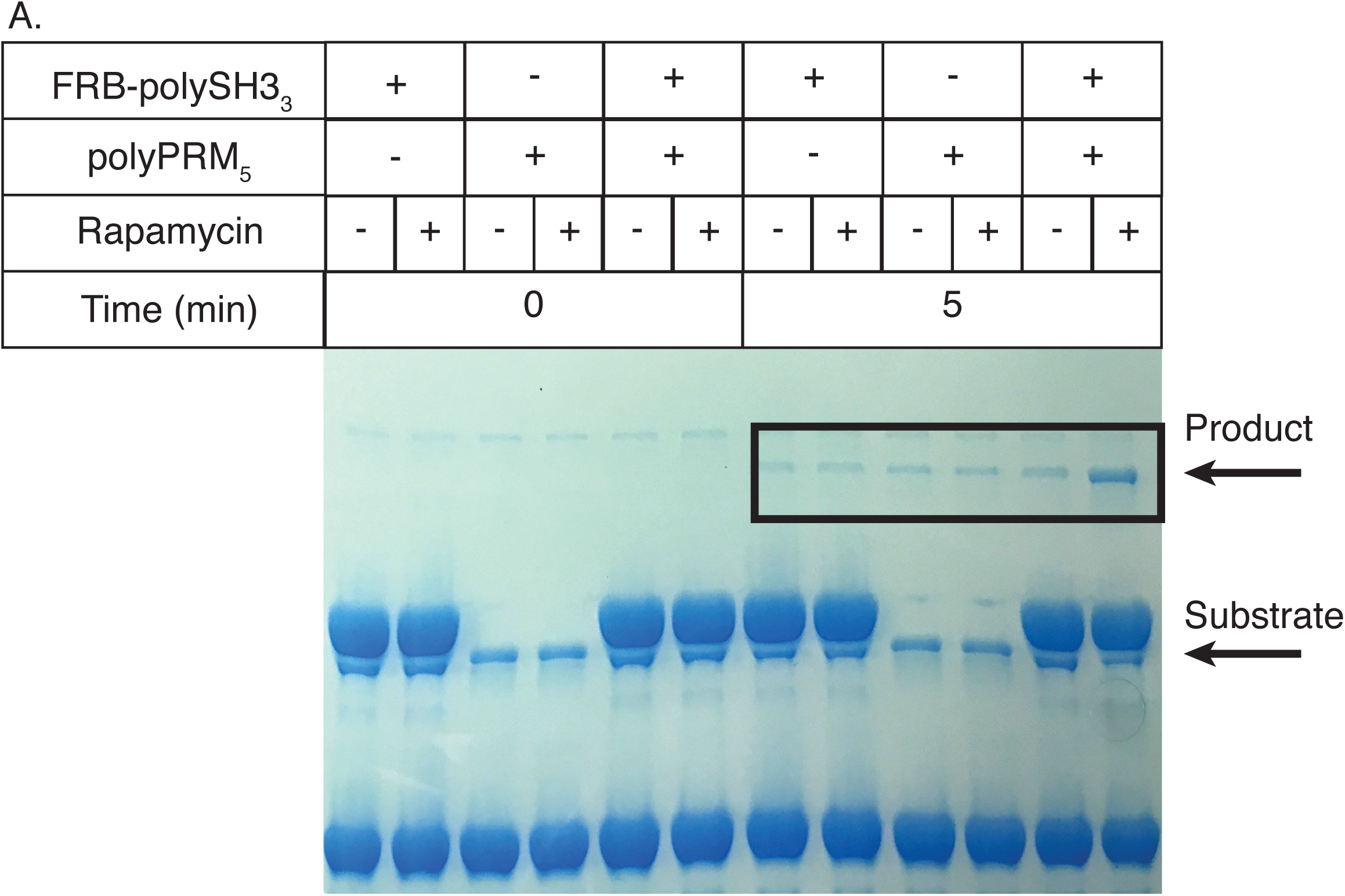
Activity enhancement requires both scaffolds to be present and both E2 and substrate to be tethered to FRB-polySH3_3_. (A) SDS-PAGE gel showing the production of SUMOylated substrate over time in the presence of FRB-polySH3_3_, polyPRM_5_ or both, with and without rapamycin.

**Figure S3.**
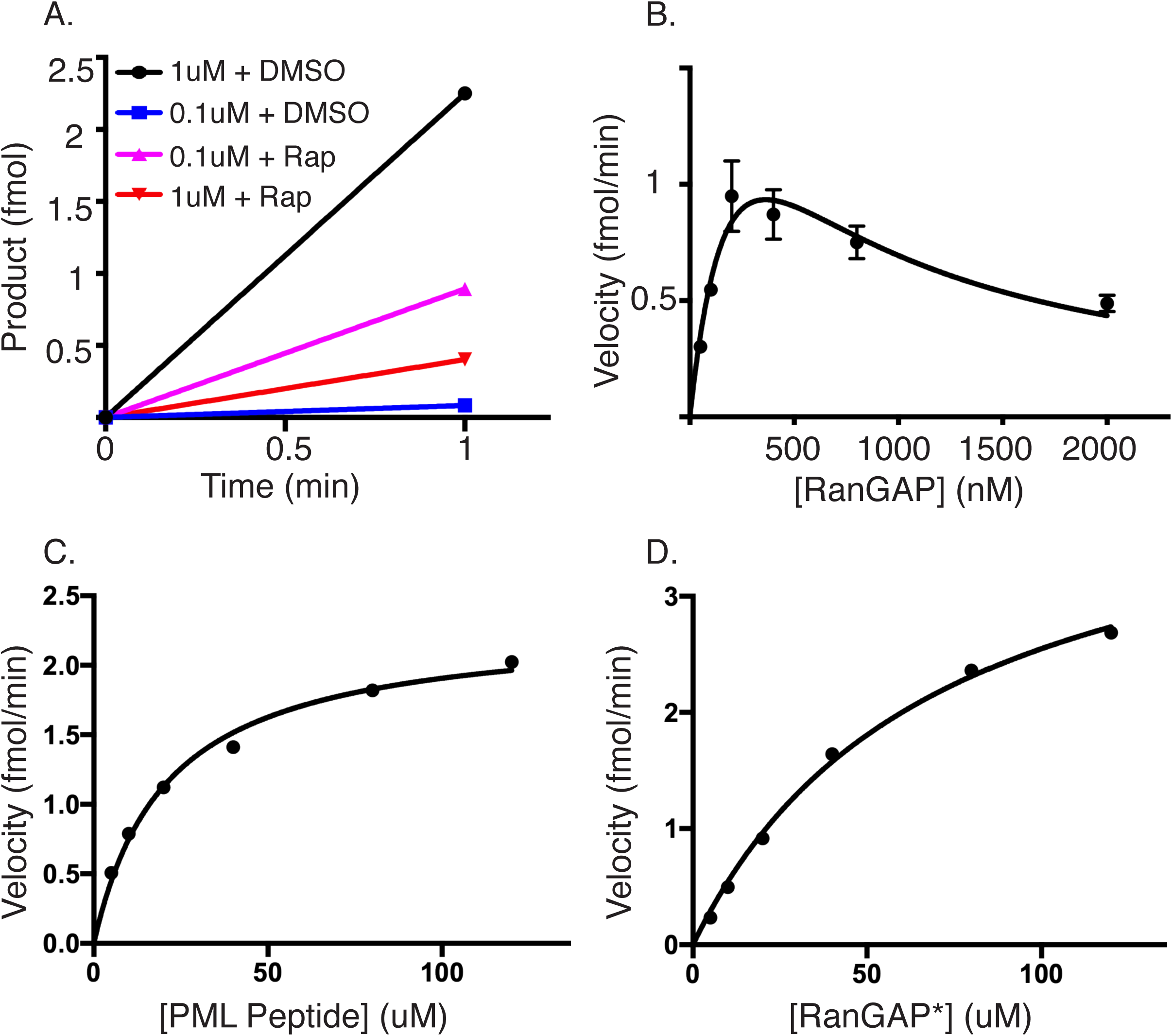
SUMOylation of RanGAP, PML peptide and RanGAP* substrates. (A) SUMOylation activity over time of RanGAP in the presence of FRB-polySH3_3_-polyPRM_5_ condensates at two different concentrations (1 and 0.1 µM) with and without rapamycin. 1uM + DMSO (black circles), 1uM + Rap (blue squares), 0.1uM + DMSO (magenta triangles), and 0.1uM + Rap (red inverted triangles). (B)-(D) SUMOylation velocity as a function of substrate concentration for RanGAP (B), PML peptide substrate (C), and RanGAP* mutant (D). RanGAP fit to substrate inhibition, while PML peptide and RanGAP* mutant fit to standard Michaelis-Menten.

**Figure S4.**
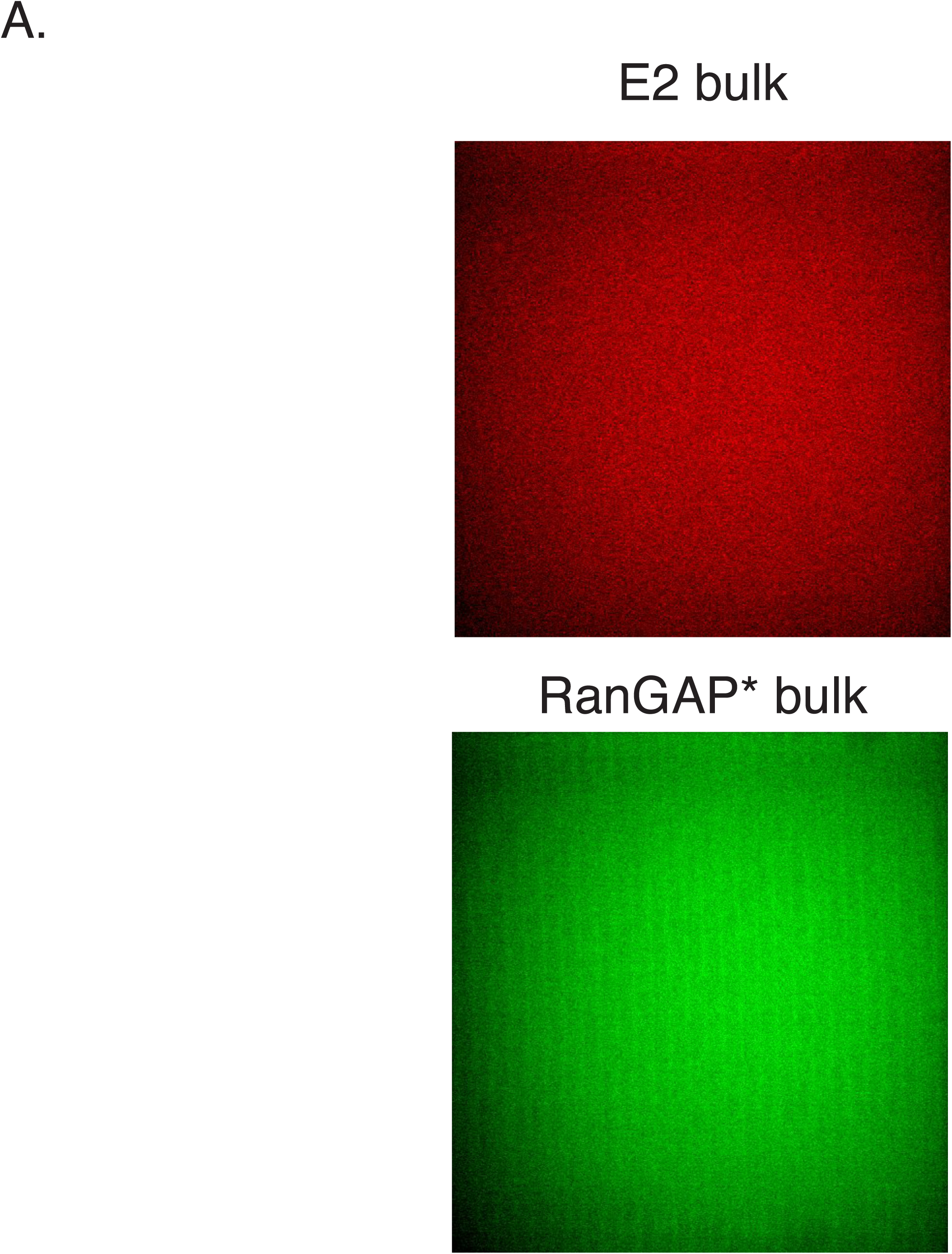
Quantitative images of bulk solution after centrifugation. Representative confocal fluorescence microscopy images of the bulk solution after clarification by centrifugation. Top row shows mCherry-FKBP-E2, bottom row shows FKBP-EGFP-RanGAP*.

**Figure S5.**
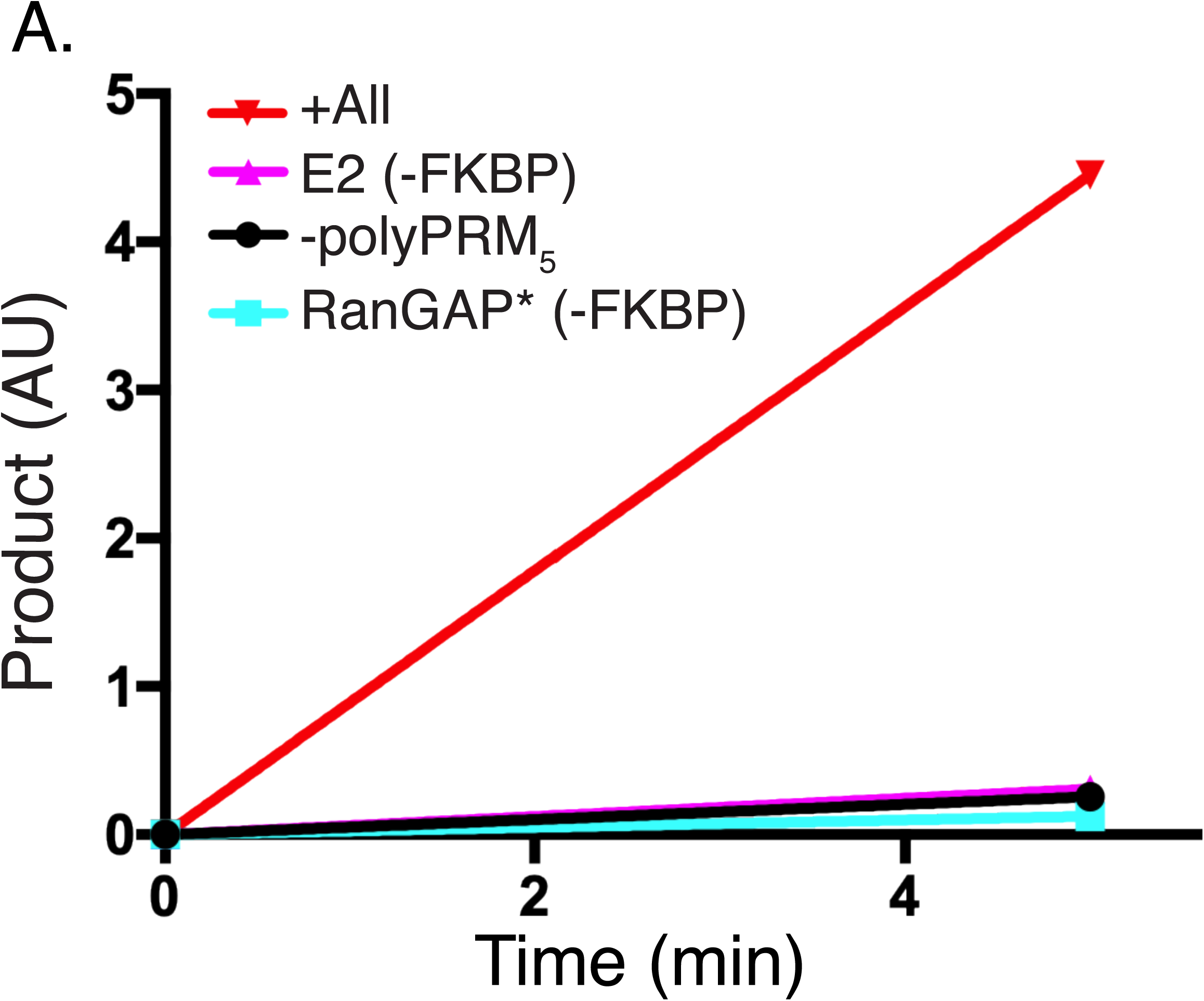
Bulk scaffold enhancement requires both scaffolds to be present and both E2 and substrate to be tethered. (A) RanGAP* SUMOylation over time at sub-critical concentrations of FRB-polySH3_3_:polyPRM_5_ without polyPRM_5_ (-polyPRM_5_, black circles), with substrate not tethered (Substrate (-FKBP), cyan squares), with E2 not tethered (E2 (-FKBP), magenta triangles), and with all components present and tethered (+All, red inverted triangles).

**Figure S6.**
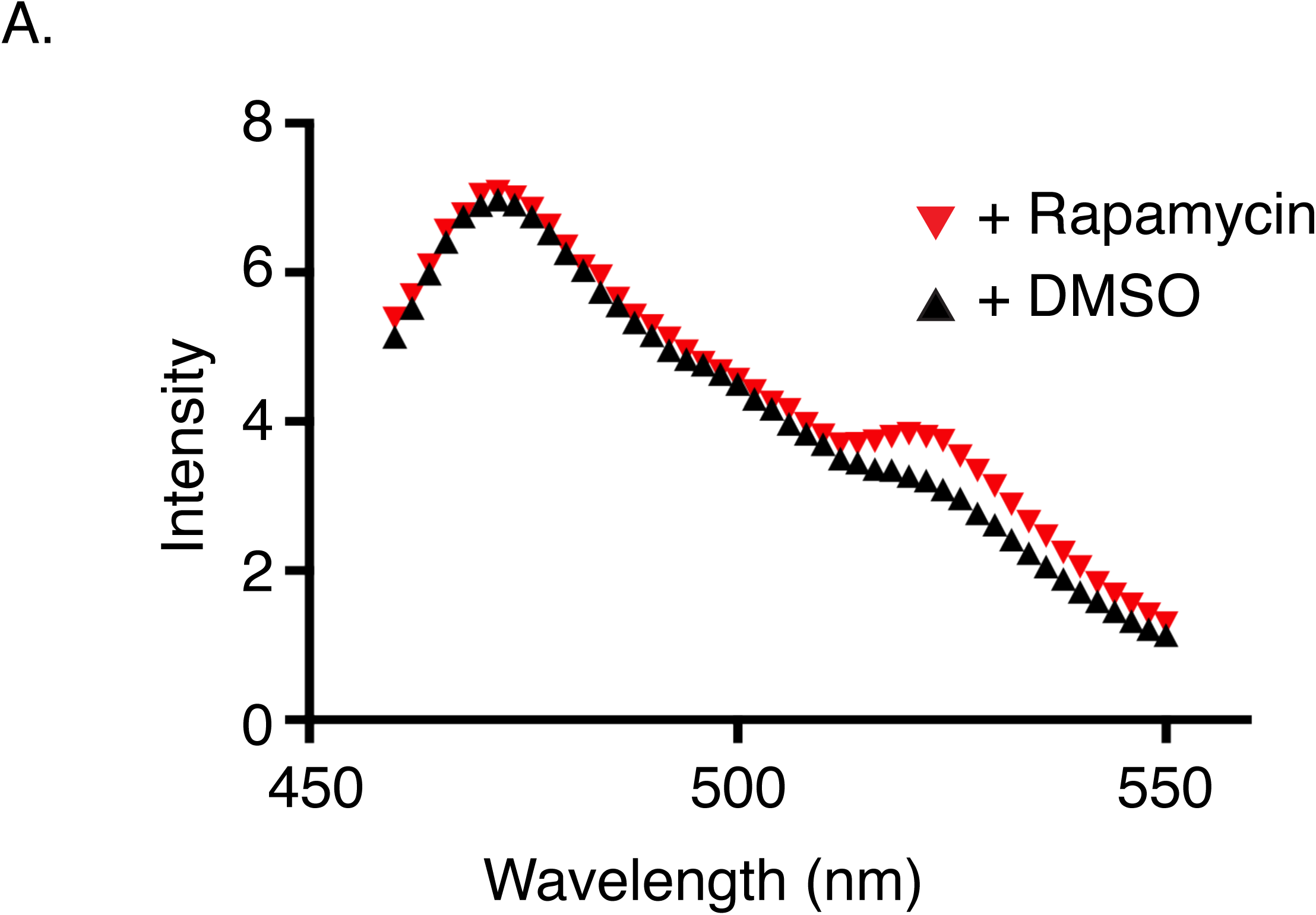
FRET increase requires rapamycin. (A) Fluorescence emission spectrum of FKBP-YPet-RanGAP* upon 445 nm excitation of CyPet-FKBP-E2. Spectra recorded in the presence of FRB-polySH3_3_ + polyPRM_5_ with either Rapamycin (inverted red triangles) or DMSO (black triangles).

**Figure S7.**
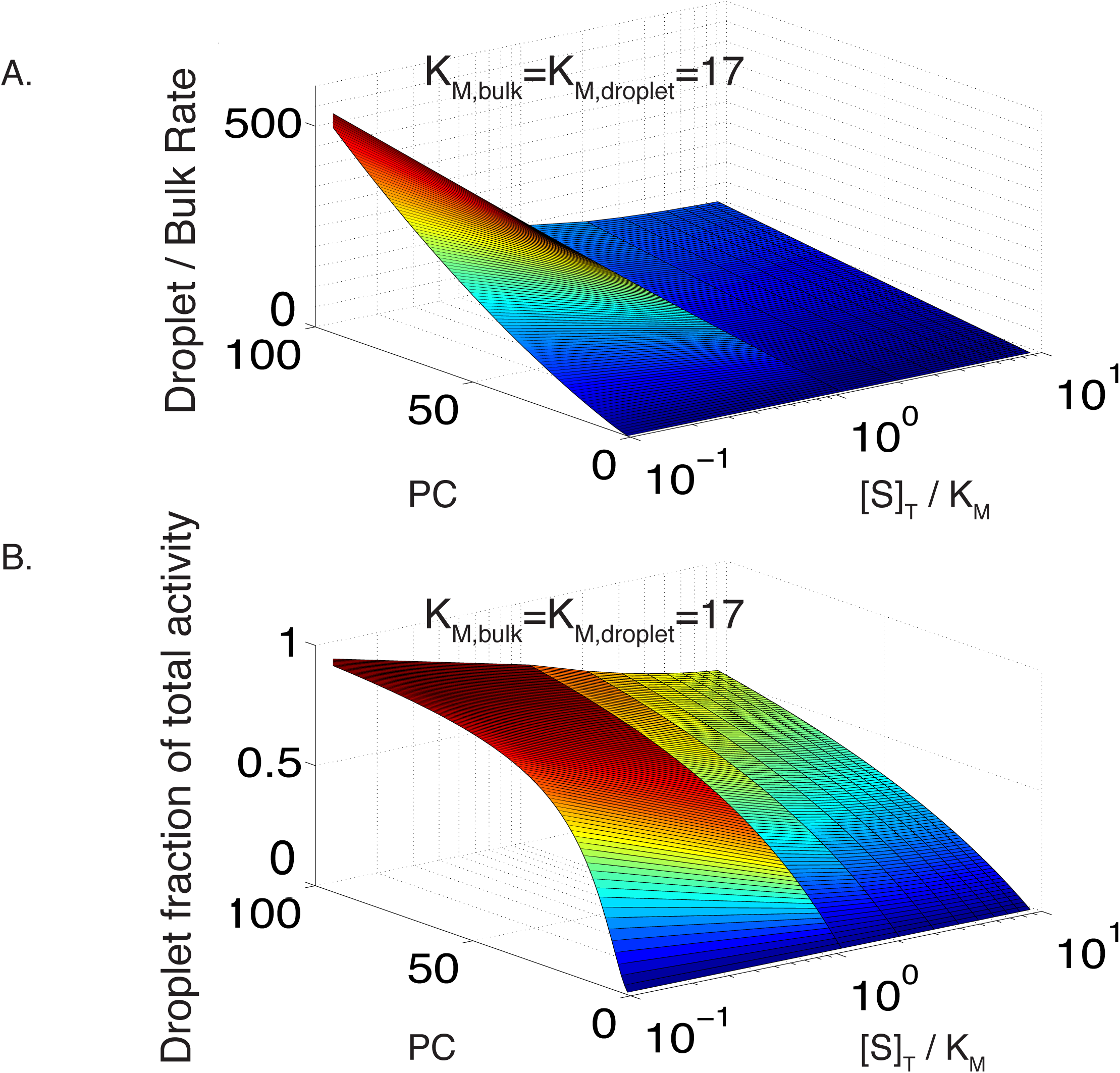
Droplet activity increases rapidly relative to bulk as a function of partition coefficient. (A) Modeled ratio of droplet and bulk reaction rates as a function of substrate concentration and partition coefficient (PC). Both reactions are scaffolded and have K_M_ = 17. Enzyme partitioning is identical to substrate partitioning, and [E] = 0.1[S]. (B) Modeled fractional activity contributed by the droplet phase as a function of substrate concentration and partition coefficient (PC). Conditions same as in (A), with a 0.01 droplet volume fraction.

**Figure S8.**
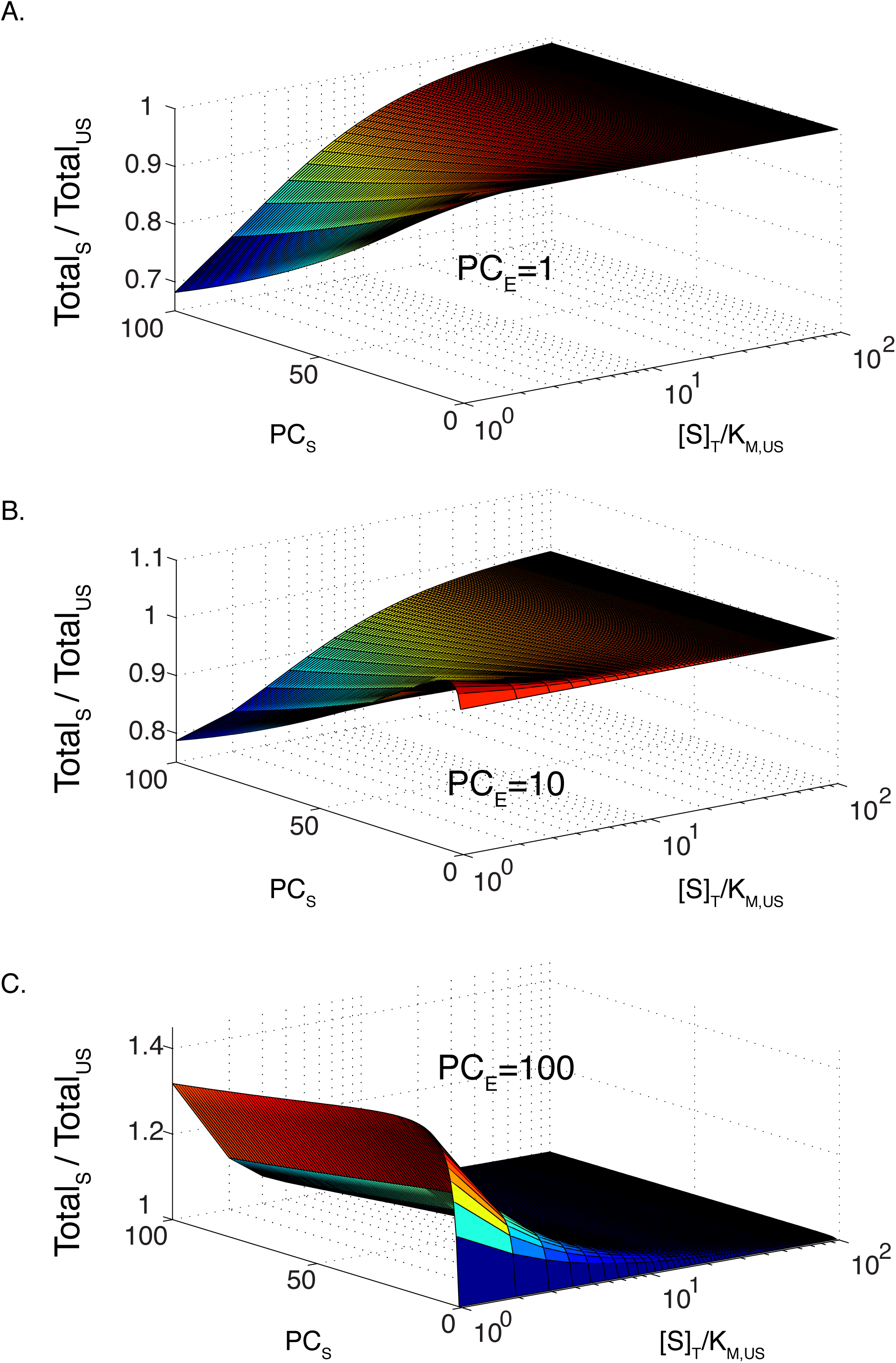
Total scaffold activity can be less than total unscaffolded activity in certain regimes if enzyme partitioning is much less than substrate partitioning. (A)-(C) Modeled ratio of total activity in a phase separated solution, with and without recruitment of enzyme and substrate to the scaffold as a function of substrate concentration and substrate partition coefficient (PC_S_). Both reactions have K_M_ = 70. Enzyme partitioning (PC_E_) is 1 (A), 10 (B), and 100 (C); enzyme concentration, [E] = 0.1[S]. Model based on 0.01 droplet volume fraction.

## Methods

### Constructs

PolyPRM_5_, polySH3_3_ and polySH3_5_ were described previously^13^. Ubc9, SAE1, and SAE2 are all full-length human proteins. SUMO1 is truncated after glycine 97 so that it is conjugation-competent^59^. The RanGAP protein used here corresponds to residues 398-587 in human RanGAP. PML peptide corresponds to residues 480-495 in human PML I. Ubc9, SAE1, SAE2, RanGAP CTD, and PML peptide substrate were each fused to FKBP12 and/or their respective fluorescent proteins using PCR. FKBP and fluorescent proteins linked together via the sequence (GGS)_4_. FRB (residues 2025 – 2115 of human mTOR) was synthesized by IDT and ligated to the N-termini of polySH3_3_ and polySH3_5_ (linked via (GGS)_4_). Exact amino acid sequence of each protein following proteolytic removal of tags (see below) is provided in Supplementary Table 1. Poly-PRM_5_, FRB-polySH3_3_, FRB-polySH3_5_, RanGAP CTD, and PML peptide constructs all contain an N-terminal MBP (maltose-binding protein) tag followed by a cleavage site for the TeV protease (ENLYFQG), followed by the insert, then another TeV cleavage site at the C-terminus, followed by a His_6_-tag. The SUMO1 construct contains an N-terminal His_8_-tag, followed by a TeV cleavage site. E2 constructs contain an N-terminal His_10_-tag, followed by a TeV cleavage site, then the insert, then another TeV cleavage site on the C-terminus, then a polybasic tag (RK)_5_. The SAE1 construct contains an N-terminal His_6_-tag followed by a TeV cleavage site. The SAE2 construct contains an N-terminal MBP-tag, followed by a TeV cleavage site.

### Protein expression and purification

All proteins were purified similarly with slight variations. All proteins except SAE1 and SAE2 were grown in *E. coli* strain BL21^TIR^ to an OD_600_ of ∼0.8 and induced with 1 mM IPTG overnight at 18 °C. To produce the E1 heterodimer, plasmids encoding SAE1 (ampicillin resistant) and SAE2 (streptomycin resistant) were co-transformed into Rosetta (DE3) bacteria (Novagen-chloramphenicol resistant), grown to OD_600_ ∼0.7, and induced with 1 mM IPTG overnight at 18 °C. For FRB-polySH3, RanGAP, PML peptide, and E1 proteins, overexpressing cells were lysed in buffer containing 50 mM Tris pH 8, 300 mM NaCl, 10 mM imidazole, 5 mM β-mercaptoethanol (BME). The lysate was cleared by centrifugation and the supernatant was applied to Ni NTA-agarose (Qiagen), which was washed with the same buffer. Proteins were eluted with buffer containing 50 mM Tris pH 8, 150 mM NaCl, 300 mM Imidazole, 5 mM BME. The eluate was loaded onto amylose resin (NEB), which was washed with 50 mM Tris pH 8, 50-150 mM NaCl, 1 mM DTT, and 1 mM EDTA (except E1). Proteins were eluted with 50 mM Tris pH 8, 50 mM NaCl, 1 mM DTT, 1 mM EDTA (except E1), and 50 mM maltose (Fisher). The amylose eluate was digested with TeV protease (∼1:100) overnight at 4 °C, filtered (0.45 µm), and loaded onto anion exchange resin (Source15Q, GE Healthcare), and eluted with a linear gradient of NaCl (50-400 mM) in 50 mM Tris pH 8, 1 mM DTT, 1 mM EDTA (except E1) buffer. Protein containing fractions were collected, filtered, and polished using gel filtration (Superdex 75 or 200, GE Healthcare) in 50 mM Tris pH 8, 150 mM NaCl, 1 mM DTT buffer. E2 and polyPRM_5_ proteins were purified using Ni NTA-agarose (Qiagen) followed by cation exchange (Source15S, GE Healthcare) chromatography, TEV digestion, and cation exchange (Source15S) and gel filtration (Superdex 75 or 200) chromatographies. Lysis and Ni-NTA wash buffers were the same as above except with 500 mM NaCl. Ni-NTA elution buffer contained 300 mM imidazole pH 7, 150 mM NaCl, 5 mM BME. Cation exchange buffers were the same as anion exchange buffers above, except used 20 mM imidazole pH 7 instead of 50 mM Tris pH 8. Gel filtration buffers were the same as above. SUMO1 was purified using Ni NTA-agarose (Qiagen), dilution, digestion with TeV, and anion exchange (Source15Q) and gel filtration (Superdex 75) chromatographies. Lysis and Ni-NTA wash buffer was 50 mM Tris pH 8, 500 mM NaCl, 20 mM Imidazole, 5 mM BME. Ni-NTA elution buffer was 50 mM Tris pH 8, 50 mM NaCl, 400 mM Imidazole, 5 mM BME. Ni-NTA eluate was diluted ∼5x into 25 mM Tris pH 8, 1 mM DTT buffer, then digested overnight with TeV (same as above). After filtration, the protein was separated by anion exchange chromatography, eluted using a 0-250 mM NaCl gradient. Gel filtration was same as above.

### *In vitro* SUMOylation assay

All reactions were and carried out in 100 µL (except bulk-see below) of 20 mM Tris, 110 mM potassium acetate, pH 7.5, 1 mM DTT at room temperature (∼22 °C). For reactions with condensates, polyPRM_5_ was added last after all other components (including rapamycin) were added to delay condensate formation until all other components were thoroughly mixed. Reactions contained 90nM E1, 100nM E2, 1uM RanGAP*, and 1uM SUMO1, 15uM FRB-polySH3_3_ (or 9uM FRB-polySH3_5_) and 9uM polyPRM_5_. Components were equilibrated for 1 hr, and then SUMOylation was initiated by addition of 1 mM ATP. Reactions included either 2 µM Rapamycin or 2% DMSO as indicated. For bulk samples, components were mixed as above, incubated at room temperature for 1 hour, centrifuged at 22 °C for 30 minutes at 21,000 g, and 50 µL was carefully transferred to a separate tube for assay. In all cases, samples were removed at the indicated timepoints (typically 0.5 – 10 minutes) and the reaction terminated by addition of an equal volume of 2X SDS PAGE loading buffer. Samples were not boiled, which allowed visualization of fluorescent proteins without stain. Samples were run on 10% SDS PAGE gels for 40 min at 240 V. Gels were either imaged directly (for fluorescent proteins) or stained with Coomassie blue and then imaged using a ChemiDoc gel imager (BioRad). Gel images were analyzed using Fiji, and band intensities (following background subtraction) were fit to extract kinetic parameters using Prism. Initial velocities were fit to either the standard Michaelis-Menten equation (V = (V_max_*[S]^h^/K_M_^h^+[S]^h^), for PML peptide, RanGAP*, and RanGAP* with FRB-polySH3_5_ + polyPRM_5_, or its allosteric variant (V = V_max_*[S]^h^/K_M_^h^+[S]^h^), for RanGAP* with FRB-polySH3_3_ + polyPRM_5_ (more appropriate equation based on the F-statistic, with α > 0.05).

### Microscopy

Corning 384-well clear bottom, untreated, assay plates were used for all microscopy experiments. Wells were treated with 5 M NaOH for 2 hours at room temperature, washed 15x with MilliQ (submerged to completely fill then emptied each time), blocked with 20% fatty acid free BSA (Fisher) for at least 2 hours at room temperature, washed 3-5 times with water and dried with argon. Samples were prepared identically to the SUMOylation reactions except without ATP. Fluorescence intensities were measured for pairs of proteins (E2 (mCherry) + substrate (EGFP), E2 (mCherry) + E1 (EGFP), and E2 (mCherry) + SUMO1 (ShadowG, a GFP variant)) to avoid spectral overlap. We analogously measured fluorescence intensities in mixtures of FRB-polySH3_3_ (0.5% FRB-polySH3_3_-EGFP), polyPRM_5_ and rapamycin. Images were acquired on a spinning disk confocal microscope (Nikon) with an EMCCD camera (Andor) and a 20x objective. Six samples were imaged and five images per sample were taken to quantify fluorescence intensity within droplets. Separately, phase separated samples were centrifuged at 22,000 g for 30 minutes at 22 °C, and 50 µL were removed, transferred to adjacent wells, and imaged to give the fluorescence intensity of the bulk. A dilution series for each protein was used to generate a standard curve, which was imaged simultaneously with the droplet and bulk samples to avoid differences in microscope performance or settings. Bulk concentrations were independently measured using the same workflow but on a fluorimeter instead of a microscope; the two approaches returned the same result (within error). At each wavelength, 15 µm z-stacks (1 µm steps) were acquired for droplet images, or a single image was acquired for all other samples. Images were analyzed in MATLAB using a custom script as described previously^18^. Briefly, the component with largest partition coefficient was used to segment droplets and bulk in the background subtracted and flat-field corrected image. The average intensity and standard deviation were extracted for the droplets (restricted to 9 – 28 µm diameter) and bulk, and converted to concentrations based on standard curves for each fluorescent protein.

### Droplet volume measurements

For direct measurements of droplet volume, we rapidly imaged a 200 µm z-stack through a droplet-containing sample using a spinning disk confocal fluorescence microscope. The images were subdivided into two 100 µm stacks. We joined each stack into a three-dimensional volume, identified droplets (3D objects counter-Fiji) and summed their volumes. Initially, small droplets were excluded by setting an intensity threshold based on the average intensity of all droplets with diameter > 2 µm. This threshold was then coupled with either a 1.3 µm or 1.0 µm diameter size threshold to identify smaller droplets. All measurements produced the same total droplet volume within error (0.9 ± 0.1 %), indicating that smaller droplets do not contribute appreciably to the volume measurement. Nevertheless, the volume determined with this approach may be somewhat lower than the true value due to undercounting of small/dim droplets. The lower stack encompassed the large droplets that had settled to the bottom of the well, while the top stack contained the small droplets that were still suspended in solution. Based on the dimensions of the well, the total solution height was approximately 5 mm, which equates to fifty 100 µm stacks. Since the second 100 µm stack represents the unsettled droplets, all the stacks except the first should be similar, so the droplet volume of the second stack was multiplied by 49 and added to the droplet volume of the first stack to give the total droplet volume in the stack. This was then divided by total volume of the stack to give the droplet volume fraction.

According to conservation of mass, the droplet volume fraction was determined from the total, droplet and bulk concentrations determined above by:

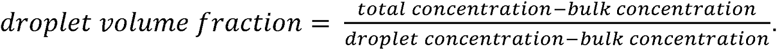

### FRET

All data were collected using a PTI fluorimeter with appropriate filters. Subcritical concentrations of the indicated scaffold mixtures were incubated with CyPet-FKBP-E2, FKBP-YPet-RanGAP* and rapamycin for 1 hr at 22 °C, centrifuged for 30 min at 21,000 g, 22 °C to remove droplets, and the supernatant was transferred to a fresh tube and imaged. Samples were excited at 445 nm and emissions collected at 460 −550 nm at 2 nm intervals. Each curve was the average of two experiments.

### Modeling

All modeling was performed in MATLAB (Mathworks). Models were generated using the Michaelis-Menten (MM) equation to describe reaction rates in the droplet and bulk compartments, R = k_cat_[E]*[S]/K_M_+[S], where [E] and [S] are the concentrations of enzyme and substrate in the respective compartment and K_M_ can either be the scaffolded (K_M,S_) or unscaffolded (K_M,US_) value. The model assumed identical k_cat_ in both compartments. The droplet and bulk concentrations are related to the total concentration according to: V_T_*C_T_= V_D_*C_D_+V_B_*C_B_, where C is concentration, V is volume, T is total, D is droplet, and B is bulk. By setting V_T_ = 1, defining the partition coefficient (PC) = C_D_/C_B_ (assumed, for simplicity, to be identical for E and S), and assigning a value to the compartment (droplet or bulk) volume fraction, this equation can be rearranged to express C_D_ or C_B_ as a function of C_T_ and PC, which can be substituted into the MM equation to yield the enzymatic rate in a given compartment.

When E and S are recruited into droplets, the droplet and bulk rates can be expressed as:

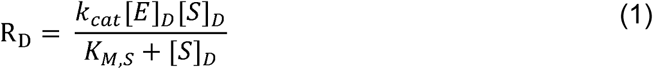

and:

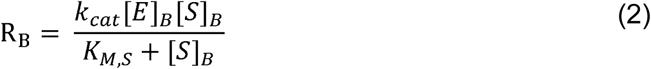

Rearranging C_T_*V_T_=V_D_*C_D_+V_B_*C_B_, and assuming a 1% droplet volume gives:

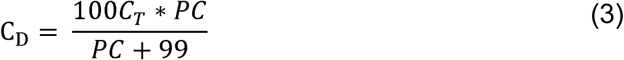

and

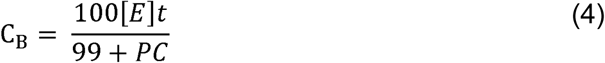

Substituting equations (3) and (4) into equations (1) and (2), respectively, gives droplet and bulk rates of:

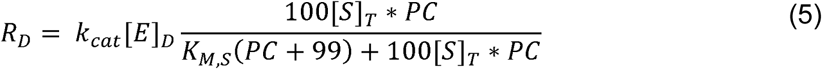

and

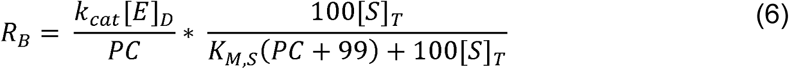

For a droplet volume of 1%, the total scaffolded reaction rate, R_T,S_ = 0.01R_D_ + 0.99 R_B_:

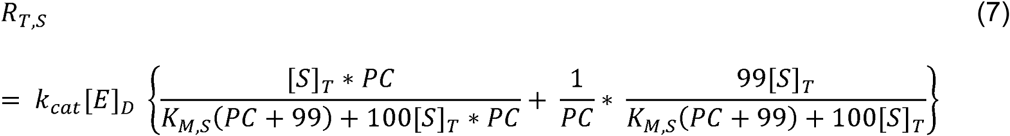

The total unscaffolded reaction rate is:

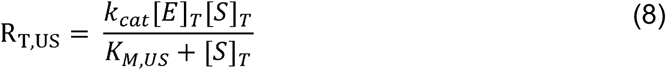

Or

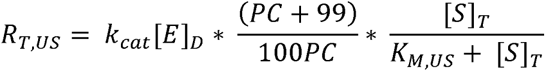

Thus, the ratio of total scaffolded to total unscaffolded rates, R_T,S_/R_T,US_ is:

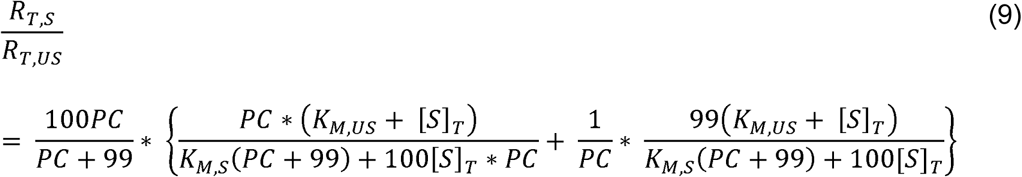

The ratio of droplet to bulk rates is given by dividing equation (5) by equation (6):

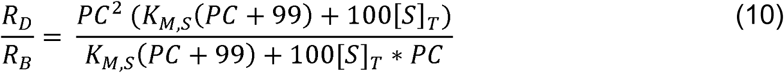

This is the ratio of rates per volume, which can be converted to the ratio of total activities by dividing by 99.

