## Supplementary material for "Phase Separation Can Increase Enzyme Activity by Concentration and Molecular Organization": Table 1

Table 1. Protein concentrations in droplets and bulk solution.

| Protein | Total Concentration (µM) | Droplet Concentration (µM) | Bulk  Concentration (µM) | Partition Coefficient |
| --- | --- | --- | --- | --- |
| E1 | 0.09 | 0.17 ± 0.01 | 0.09 ± 0.01 | 1.9 ± 0.2 |
| E2 | 0.1 | 1.4 ± 0.1 | 0.09 ± 0.01 | 14 ± 2 |
| Substrate | 1.0 | 31 ± 2 | 0.65 ± 0.03 | 48 ± 4 |
| SUMO1 | 1.0 | 1.0 ± 0.1 | 1.0 ± 0.1 | 1.0 ± 0.1 |
| FRB-polySH3_5_ | 15.0 | 660 ± 16 | 6.3 ± 0.3 | 105 ± 6 |
