## Supplemental Table 1 for "Phase Separation Can Increase Enzyme Activity by Concentration and Molecular Organization"

Supplementary Table 1. Constructs used in this study. Protein sequences after proteolytic removal of affinity tags are shown.

| Construct | Protein sequence | Tag(s) |
| --- | --- | --- |
| polyPRM_5_ | GHMKGGSWGGSKKKKTAPTPPKRSGGSGGSGGSGGSKKKKTAPTPPKRSGGSGGSGGSGGSKKKKTAPTPPKRSGGSGGSGGSGGSKKKKTAPTPPKRSGGSGGSGGSGGSKKKKTAPTPPKRSGGSGSENLYFQ | N-terminal MBP (maltose binding protein)  C-terminal His_6_ |
| FRB-polySH3_3_ | GEFMLEMWHEGLEEASRLYFGERNVKGMFEVLEPLHAMMERGPQTLKETSFNQAYGRDLMEAQEWCRKYMKSGNVKDLTQAWDLYYHVFRRISKQVDGGSGGSGGSGGSHMDLNMPAYVKFNYMAEREDELSLIKGTKVIVMEKSSDGWWRGSYNGQVGWFPSNYVTEEGDSPLASGAGGSEGGGSEGGTSGATDLNMPAYVKFNYMAEREDELSLIKGTKVIVMEKSSDGWWRGSYNGQVGWFPSNYVTEEGDSPLASGAGGSEGGGSEGGTSGATDLNMPAYVKFNYMAEREDELSLIKGTKVIVMEKSSDGWWRGSYNGQVGWFPSNYVTEEGDSPLGGGSENLYFQ | N-terminal MBP  C-terminal His_6_ |
| FRB-polySH3_5_ | GEFMLEMWHEGLEEASRLYFGERNVKGMFEVLEPLHAMMERGPQTLKETSFNQAYGRDLMEAQEWCRKYMKSGNVKDLTQAWDLYYHVFRRISKQVDGGSGGSGGSGGSKDAQTNSSSNNNNNNNNNNLGIEGRISHMDLNMPAYVKFNYMAEREDELSLIKGTKVIVMEKSSDGWWRGSYNGQVGWFPSNYVTEEGDSPLASGAGGSEGGGSEGGTSGATDLNMPAYVKFNYMAEREDELSLIKGTKVIVMEKSSDGWWRGSYNGQVGWFPSNYVTEEGDSPLASGAGGSEGGGSEGGTSGATHMDLNMPAYVKFNYMAEREDELSLIKGTKVIVMEKSSDGWWRGSYNGQVGWFPSNYVTEEGDSPLASGAGGSEGGGSEGGTSGATDLNMPAYVKFNYMAEREDELSLIKGTKVIVMEKSSDGWWRGSYNGQVGWFPSNYVTEEGDSPLASGAGGSEGGGSEGGTSGATDLNMPAYVKFNYMAEREDELSLIKGTKVIVMEKSSDGWWRGSYNGQVGWFPSNYVTEEGDSPLGGGSENLYFQ | N-terminal MBP  C-terminal His_6_ |
| SUMO1 | GEFVSKGEELFTGVVPILVELDGDVNGHKFSVSGEGEGDATYGKLTLKLICTTGKLPVPWPTLVTTFGYGLMCFARYPDHMKQHDFFKSAMPEGYVQERTIFFKDDGNYKTRAEVKFEGDTLVNRIELKGIDFKEDGNILGHKLEYNWNSHNVYIMADKQKNGIKVNFKIRHNIEDGSVQLADHYQQNTPIGDGPVLLPDNHYLSTQSKLSKDPNEKRDHMVLLEFVTAAGITLGMDELYKSGLRSRAQASNSAVDGTMSDQEAKPSTEDLGDKKEGEYIKLKVIGQDSSEIHFKVKMTTHLKKLKESYCQRQGVPMNSLRFLFEGQRIADNHTPKELGMEEEDVIEVYQEQTGG | N-terminal His_8_ |
| SAE1 | GEFGGSGGSGGSGGSMVEKEEAGGGISEEEAAQYDRQIRLWGLEAQKRLRASRVLLVGLKGLGAEIAKNLILAGVKGLTMLDHEQVTPEDPGAQFLIRTGSVGRNRAEASLERAQNLNPMVDVKVDTEDIEKKPESFFTQFDAVCLTCCSRDVIVKVDQICHKNSIKFFTGDVFGYHGYTFANLGEHEFVEEKTKVAKVSQGVEDGPGPDTKRAKLDSSETTMVKKKVVFCPVKEALEVDWSSEKAKAALKRTTSDYFLLQVLLKFRTDKGRDPSSDTYEEDSELLLQIRNDVLDSLGISPDLLPEDFVRYCFSEMAPVCAVVGGILAQEIVKALSQRDPPHNNFFFFDGMKGNGIVECLGPK | N-terminal His_6_ |
| SAE2-mGarnet | GEFSGGSGGSGGSMALSRGLPRELAEAVAGGRVLVVGAGGIGCELLKNLVLTGFSHIDLIDLDTIDVSNLNRQFLFQKKHVGRSKAQVAKESVLQFYPKANIVAYHDSIMNPDYNVEFFRQFILVMNALDNRAARNHVNRMCLAADVPLIESGTAGYLGQVTTIKKGVTECYECHPKPTQRTFPGCTIRNTPSEPIHCIVWAKYLFNQLFGEEDADQEVSPDRADPEAAWEPTEAEARARASNEDGDIKRISTKEWAKSTGYDPVKLFTKLFKDDIRYLLTMDKLWRKRKPPVPLDWAEVQSQGEETNASDQQNEPQLGLKDQQVLDVKSYARLFSKSIETLRVHLAEKGDGAELIWDKDDPSAMDFVTSAANLRMHIFSMNMKSRFDIKSMAGNIIPAIATTNAVIAGLIVLEGLKILSGKIDQCRTIFLNKQPNPRKKLLVPCALDPPNPNCYVCASKPEVTVRLNVHKVTVLTLQDKIVKEKFAMVAPDVQIEDGKGTILISSEEGETEANNHKKLSEFGIRNGSRLQADDFLQDYTLLINILHSEDLGKDVEFEVVGDAPEKVGPKQAEDAAKSITNGSDDGAQPSTSTAQEQDDVLIVDSDEEDSSNNADVSEEERSRKRKLDEKENLSAKRSRIEQKEELDDVIALDLDLRSRAQASNSAVDGTNSLIKENMRMKVVLEGSVNGHQFKCTGEGEGNPYMGTQTMRIKVIEGGPLPFAFDILATSFMYGSKTFIKYPKGIPDFFKQSFPEGFTWERVTRYEDGGVITVMQDTSLEDGCLVYHAQVRGVNFPSNGAVMQKKTKGWEPNTEMMYPADGGLRGYNHMALKVDGGGHLSCSLVTTYRSKKTVGNIKMPGIHAVDRRLERLEESDNEMFVVQREHAVAKFAGLGGG | N-terminal-MBP |
| mCherry-E2 | GEFVSKGEEDNMAIIKEFMRFKVHMEGSVNGHEFEIEGEGEGRPYEGTQTAKLKVTKGGPLPFAWDILSPQFMYGSKAYVKHPADIPDYLKLSFPEGFKWERVMNFEDGGVVTVTQDSSLQDGEFIYKVKLRGTNFPSDGPVMQKKTMGWEASSERMYPEDGALKGEIKQRLKLKDGGHYDAEVKTTYKAKKPVQLPGAYNVNIKLDITSHNEDCTIVEQYERAEGRHSTGGMDELYKSGGGSGGSGGSGGSMSGIALSRLAQERKAWRKDHPFGFVAVPTKNPDGTMNLMNWECAIPGKKGTPWEGGLFKIRMLFKDDYPSSPPKCKFEPPLFHPNVYPSGTVCLSILEEDKDWRPAITIKQILLGIQELLNEPNIQDPAQAEAYTIYCQNRVEYEKRVRAQAKKFAPSLEENLYFQ | N-terminal His_10_  C-terminal RK5  (5 repeats of arginine and lysine) |
| mCherry-FKBP-E2 | GEFVSKGEEDNMAIIKEFMRFKVHMEGSVNGHEFEIEGEGEGRPYEGTQTAKLKVTKGGPLPFAWDILSPQFMYGSKAYVKHPADIPDYLKLSFPEGFKWERVMNFEDGGVVTVTQDSSLQDGEFIYKVKLRGTNFPSDGPVMQKKTMGWEASSERMYPEDGALKGEIKQRLKLKDGGHYDAEVKTTYKAKKPVQLPGAYNVNIKLDITSHNEDCTIVEQYERAEGRHSTGGMDELYKQLMGVQVETISPGDGRTFPKRGQTCVVHYTGMLEDGKKFDSSRDRNKPFKFMLGKQEVIRGWEEGVAQMSVGQRAKLTISPDYAYGATGHPGIIPPHATLVFDVELLKLNEGGSGGSGGSGGSLRSRAQASNSAVDGTMSGIALSRLAQERKAWRKDHPFGFVAVPTKNPDGTMNLMNWECAIPGKKGTPWEGGLFKLRMLFKDDYPSSPPKCKFEPPLFHPNVYPSGTVCLSILEEDKDWRPAITIKQILLGIQELLNEPNIQDPAQAEAYTIYCQNRVEYEKRVRAQAKKFAPSLENLYFQ | N-terminal His_10_  C-terminal RK5 |
| CyPet-FKBP-E2 | GEFVSKGEELFGGIVPILVELEGDVNGHKFSVSGEGEGDATYGKLTLKFICTTGKLPVPWPTLVTTLTWGVQCFSRYPDHMKQHDFFKSVMPEGYVQERTIFFKDDGNYKTRAEVKFEGDTLVNRIELKGIDFKEDGNILGHKLEYNYISHNVYITADKQKNGIKANFKARHNITDGSVQLADHYQQNTPIGDGPVILPDNHYLSTQSALSKDPNEKRDHMVLLEFVTAAGITHGMDELYKQLMGVQVETISPGDGRTFPKRGQTCVVHYTGMLEDGKKFDSSRDRNKPFKFMLGKQEVIRGWEEGVAQMSVGQRAKLTISPDYAYGATGHPGIIPPHATLVFDVELLKLNEGGSGGSGGSGGSLRSRAQASNSAVDGTMSGIALSRLAQERKAWRKDHPFGFVAVPTKNPDGTMNLMNWECAIPGKKGTPWEGGLFKLRMLFKDDYPSSPPKCKFEPPLFHPNVYPSGTVCLSILEEDKDWRPAITIKQILLGIQELLNEPNIQDPAQAEAYTIYCQNRVEYEKRVRAQAKKFAPSLENLYFQ | N-terminal His_10_  C-terminal RK5 |
| FKBP-EGFP-RanGAP | GEFMGVQVETISPGDGRTFPKRGQTCVVHYTGMLEDGKKFDSSRDRNKPFKFMLGKQEVIRGWEEGVAQMSVGQRAKLTISPDYAYGATGHPGIIPPHATLVFDVELLKLNEGGSGGSGGSGGSVSKGEELFTGVVPILVELDGDVNGHKFSVSGEGEGDATYGKLTLKFICTTGKLPVPWPTLVTTLTYGVQCFSRYPDHMKQHDFFKSAMPEGYVQERTIFFKDDGNYKTRAEVKFEGDTLVNRIELKGIDFKEDGNILGHKLEYNYNSHNVYIMADKQKNGIKVNFKIRHNIEDGSVQLADHYQQNTPIGDGPVLLPDNHYLSTQSKLSKDPNEKRDHMVLLEFVTAAGITLGMDELYKSGLRSPQQRGQGEKSATPSRKILDPNTGEPAPVLSSPPPADVSTFLAFPSPEKLLRLGPKSSVLIAQQTDTSDPEKVVSAFLKVSSVFKDEATVRMAVQDAVDALMQKAFNSSSFNSNTFLTRLLVHMGLLKSEDKVKAIANLYGPLMALNHMVQQDYFPKALAPLLLAFVTKPNSALESCSFARHSLLQTLSKVGSENLYFQ | N-terminal MBP  C-terminal His_6_ |
| FKBP-EGFP-RanGAP* | GEFMGVQVETISPGDGRTFPKRGQTCVVHYTGMLEDGKKFDSSRDRNKPFKFMLGKQEVIRGWEEGVAQMSVGQRAKLTISPDYAYGATGHPGIIPPHATLVFDVELLKLNEGGSGGSGGSGGSVSKGEELFTGVVPILVELDGDVNGHKFSVSGEGEGDATYGKLTLKFICTTGKLPVPWPTLVTTLTYGVQCFSRYPDHMKQHDFFKSAMPEGYVQERTIFFKDDGNYKTRAEVKFEGDTLVNRIELKGIDFKEDGNILGHKLEYNYNSHNVYIMADKQKNGIKVNFKIRHNIEDGSVQLADHYQQNTPIGDGPVLLPDNHYLSTQSKLSKDPNEKRDHMVLLEFVTAAGITLGMDELYKSGLRSPQQRGQGEKSATPSRKILDPNTGEPAPVLSSPPPADVSTFLAFPSPEKLLRLGPKSSVLIAQQTDTSDPEKVVSAFLKVSSVFKDEATVRMAVQDAVDALMQKAFNSSSFNSNTFLTRLLVHMGLLKSEDKVKAIANLYGPLMALNHMVQQDYFPKALAPLLLAAVTKPNSALESCSFARHSLLQTLSKVGSENLYFQ | N-terminal MBP  C-terminal His_6_  Mutation site colored red |
| FKBP-YPet-RanGAP* | GEFMGVQVETISPGDGRTFPKRGQTCVVHYTGMLEDGKKFDSSRDRNKPFKFMLGKQEVIRGWEEGVAQMSVGQRAKLTISPDYAYGATGHPGIIPPHATLVFDVELLKLNEGGSGGSGGSGGSVSKGEELFTGVVPILVELDGDVNGHKFSVSGEGEGDATYGKLTLKLLCTTGKLPVPWPTLVTTLGYGVQCFARYPDHMKQHDFFKSAMPEGYVQERTIFFKDDGNYKTRAEVKFEGDTLVNRIELKGIDFKEDGNILGHKLEYNYNSHNVYITADKQKNGIKANFKIRHNIEDGGVQLADHYQQNTPIGDGPVLLPDNHYLSYQSALFKDPNEKRDHMVLLEFLTAAGITEGMNELYKSGLRSPQQRGQGEKSATPSRKILDPNTGEPAPVLSSPPPADVSTFLAFPSPEKLLRLGPKSSVLIAQQTDTSDPEKVVSAFLKVSSVFKDEATVRMAVQDAVDALMQKAFNSSSFNSNTFLTRLLVHMGLLKSEDKVKAIANLYGPLMALNHMVQQDYFPKALAPLLLAAVTKPNSALESCSFARHSLLQTLSKVGSENLYFQ | N-terminal MBP  C-terminal His_6_  Mutation site colored red |
| FKBP-EGFP-PML peptide | GEFMGVQVETISPGDGRTFPKRGQTCVVHYTGMLEDGKKFDSSRDRNKPFKFMLGKQEVIRGWEEGVAQMSVGQRAKLTISPDYAYGATGHPGIIPPHATLVFDVELLKLNEGGSGGSGGSGGSVSKGEELFTGVVPILVELDGDVNGHKFSVSGEGEGDATYGKLTLKFICTTGKLPVPWPTLVTTLTYGVQCFSRYPDHMKQHDFFKSAMPEGYVQERTIFFKDDGNYKTRAEVKFEGDTLVNRIELKGIDFKEDGNILGHKLEYNYNSHNVYIMADKQKNGIKVNFKIRHNIEDGSVQLADHYQQNTPIGDGPVLLPDNHYLSTQSKLSKDPNEKRDHMVLLEFVTAAGITLGMDELYKGGSGGSGGSKVDVIDLTIESSSDEEEDPPAKRGSAGSAGSAGSAGSAGSAGSAGSAGSAGSAGSAGSASQTQSPRKVIKMESEEGSENLYFQ | N-terminal MBP  C-terminal His_6_ |
| EGFP-PML peptide | GEFVSKGEELFTGVVPILVELDGDVNGHKFSVSGEGEGDATYGKLTLKFICTTGKLPVPWPTLVTTLTYGVQCFSRYPDHMKQHDFFKSAMPEGYVQERTIFFKDDGNYKTRAEVKFEGDTLVNRIELKGIDFKEDGNILGHKLEYNYNSHNVYIMADKQKNGIKVNFKIRHNIEDGSVQLADHYQQNTPIGDGPVLLPDNHYLSTQSKLSKDPNEKRDHMVLLEFVTAAGITLGMDELYKGGSGGSGGSKVDVIDLTIESSSDEEEDPPAKRGSAGSAGSAGSAGSAGSAGSAGSAGSAGSAGSAGSASQTQSPRKVIKMESEEGSENLYFQ | N-terminal MBP  C-terminal His_6_ |
